# Controls of SAR11 subclade abundance, diversity, and growth in two Mid-Atlantic estuaries

**DOI:** 10.1101/2022.05.04.490708

**Authors:** Barbara J. Campbell, Shen Jean Lim, David L. Kirchman

## Abstract

SAR11 is a dominant bacterial clade in marine oligotrophic ecosystems. SAR11 can also be dominant in estuarine systems, where they are not well-studied. We examined the effects of season, nutrient concentrations, and salinity in shaping SAR11 subclade abundance, diversity, function, and growth in two Mid-Atlantic estuaries, the Delaware and Chesapeake Bays. Using metagenome-assembled genomes, we identified twelve distinct genomospecies within the Ia, II, IIIa, and V subclades, which made up to 60% of the total bacterial community. The functional potential of all SAR11 genomospecies varied, especially in carbohydrate metabolism, transporters, and one-carbon metabolic pathways. Predicted growth rates, estimated by the Peak to Trough method, varied by season and genomospecies. SAR11 growth rates negatively correlated in the spring but positively correlated in the summer with chlorophyll *a* concentrations and bacterial production, as well as phosphate and ammonium concentrations. Genomospecies in Ia.1, IIIa.2, and IIIa.4 subclades had low growth rates, while genomospecies in Ia.3, Ia.5, Ia.6, II, and V subclades had higher and more variable growth rates that were positively correlated with phosphate concentrations and temperature. Growth rate variation between subclades was associated with carbohydrate metabolic gene repertoires, especially glycolysis and number of transporters. While total transcript to genome ratios generally mirrored growth rates, transcription of genes involved in phosphate and nitrogen transport were negatively associated with growth rates. These data suggest that SAR11 genomospecies abundance varies in these estuaries because of differences in growth rates and metabolic capacities in response to changes in environmental conditions.

**Importance:** The SAR11 clade is one of the most abundant bacterial groups in marine systems, including many estuaries. From the Delaware Bay and Chesapeake Bay environmental metagenomes, we reconstructed nearly complete SAR11 metagenome-assembled genomes representing ten genomospecies in four subclades, of which at least one is novel. Growth rate estimates of genomospecies correlated with functional gene repertoires of carbohydrate transporter and metabolism. Different SAR11 genomospecies dominated among the seasons, depending on their growth rates, biological productivity, and nutrient concentrations. Our RNAseq approach facilitated an understanding of the environmental controls on the abundance of SAR11 genomospecies in their natural habitat. This study is the first to combine multiple measures of diversity, abundance, functional potential, growth rates and activity of this important group, demonstrating a direct link between SAR11 genomospecies abundance and growth in the context of its environment.

## Introduction

The SAR11 clade of bacteria (*Pelagibacterales*) represents about 25% of all marine planktonic cells in the stratified, oligotrophic open ocean (1, 2). The clade comprises at least nine oceanic subclades (Ia.1, Ia.3, Ib, Ic, IIa, IIb, IIIa, IV and V) and one freshwater subclade (IIIb) (3–5). Although typically less than 5% of the microbial community in freshwater systems, SAR11 members unexpectedly may comprise up to 60% of free-living bacteria in estuaries (6). Estuaries typically are hot spots of production with nutrient concentrations ten-fold higher or more than the open ocean. Various SAR11 subclades such as Ia.1, II and IIIa are found in estuaries including the Delaware Bay and some taxa are estuary-specific (2, 7–10). Nevertheless, SAR11 subclade dominance in estuaries is not universal and may depend on season and salinity (6, 7, 9, 11).

SAR11 are aerobic heterotrophs with small, streamlined genomes that are diverse across subclades (12, 13). Analyses of seven published genomes in four marine SAR11 subclades (Ia.1, Ia.3, IIIa, V) revealed common genomic features, including low G+C content, low numbers of paralogs, a large hypervariable region, and high gene synteny, as well as core genes that comprised about 50% of their genomes (13). Transport genes make up approximately 15% of these genomes, which is around twice that of other bacteria, including the marine *Roseobacter* clade from the same alphaproteobacterial class (13–15). Core SAR11 genes encode ATP-binding cassette (ABC) transport systems for amino acids, iron transport, and tripartite ATP-independent periplasmic (TRAP) transporter(s) for C4 compounds. Other common genes participate in photoheterotrophy (13, 16), electron transport, and the TCA cycle (13).

Despite their commonalities, several important distinctions exist among SAR11 subclades. Their abundance differs among geographic locations (2, 5) and with different environmental parameters, including light (17–19), temperature, and nutrient concentrations, which also vary with depth (17, 18, 20, 21). Accordingly, SAR11 subclades and genomes differ in the presence of ecologically important genes, such as those involved in the transport of organic carbon, sulfur, and phosphorus, as well as the oxidation of carbon (e.g., C1) compounds (13, 22). However, the number of sequenced genomes used to make these observations is low compared to the abundance and diversity of the SAR11 clade. There is little known about the genomic makeup of several subclades, as well as SAR11 subclade distribution in estuaries.

In addition to the lack of information about SAR11 subclade ecogenomics in estuaries, the differences in SAR11 activity or growth between subclades is not well studied. Some of this is because of the paucity of cultured SAR11 representatives (3). The slow growth rate of *P. ubique* and relatives in the laboratory, averaging about 0.4 per day (12, 23–25), raises questions about growth rates and activity of this group in the ocean. Previous estimates by our group using 16S rRNA gene:rRNA ratios in the coastal Mid-Atlantic suggest that about half of the SAR11 taxa had lower growth rates than average, whereas the other half had average or above average growth rates, which varied by season (26). Differential use of glucose and phosphate was also observed within the SAR11 clade, likely contributing to differences in ocean productivity (27, 28) and perhaps also to growth rate (29). In addition, data from a technique combining microautoradiography and fluorescence in situ hybridization (MAR-FISH) suggest that SAR11 bacteria are in fact growing as fast, if not faster than other bacteria in surface waters of the North Atlantic (30, 31). However, oceanographic studies found that global SAR11 gene transcription was generally lower than expected based on abundance (32) and varied in response to light and the activity of other microbes in oligotrophic open oceans (33–35).

Estuaries are ideal environments to study the abundance, activity, and functional dynamics of SAR11 subclades since these ecosystems predictably vary in time or space. The goal of this study was to determine SAR11 taxonomic, genomic, and functional diversity as well as growth and activity along environmental gradients in two contrasting but geographically close estuaries, the Delaware and Chesapeake Bays. The Delaware Bay has a steep salinity gradient, is well-mixed with little to no stratification. Primary production is limited by light in the winter, phosphorus in the late spring, and nitrogen in the summer (36). In contrast, the Chesapeake Bay is highly stratified, especially in the summer, has higher primary productivity, and contains much more mesohaline waters because it is less influenced by tides due to its length (37–39). Similar to the Delaware Bay, primary productivity in the Chesapeake varies by light, phosphorus, and nitrogen, depending on the season and location within the estuary (40). We hypothesized that these environmental gradients would structure and influence the abundance, taxonomic, and functional variability of SAR11 subclades as well as their growth and activity. To test this hypothesis, we utilized medium and high-quality assembled MAGs and reference genomes to examine the taxonomic relationships, abundances, and functional potentials of the *Pelagibacterales* in these estuaries. We also investigated differences in genomospecies growth and activity analyzing differences in metagenomes (MG) and metatranscriptomes (MT) between environmental conditions.

## Results

### MAG characteristics

Forty-four MAGs classified as *Pelagibacterales* were identified from 36 metagenome (MG) samples from five cruises to the Delaware or Chesapeake Bays in 2014 and 2015 (41) (**Table S1**). They were, on average, 88.6% complete with 0.4% redundancy. Only one MAG contained partial 16S or 23S rRNA genes. Based on phylogenomic and AAI analyses, these MAGs clustered into four larger clades (Ia, II, IIIa and V) with other known *Pelagibacterales* genomes, SAGs, or MAGs (**Fig. 1, see Fig. S1 at 10.6084/m9.figshare.19701379**). Within Ia, five different genomospecies (Ia.1, Ia.3, Ia.4, Ia.,5, and Ia.6) were identified. Most genomospecies had representative MAGs in the Delaware and Chesapeake Bays. We did not recover MAGs related to 1a.4, Ib, Ic, IIa.B or IIIb from the two bays. Novel to this study are well-represented genomospecies from II and Ia.6, as well as a much-expanded IIIa (genomospecies IIIa.4). The subclades are examined in more detail in the supplemental text.

**Figure 1.**
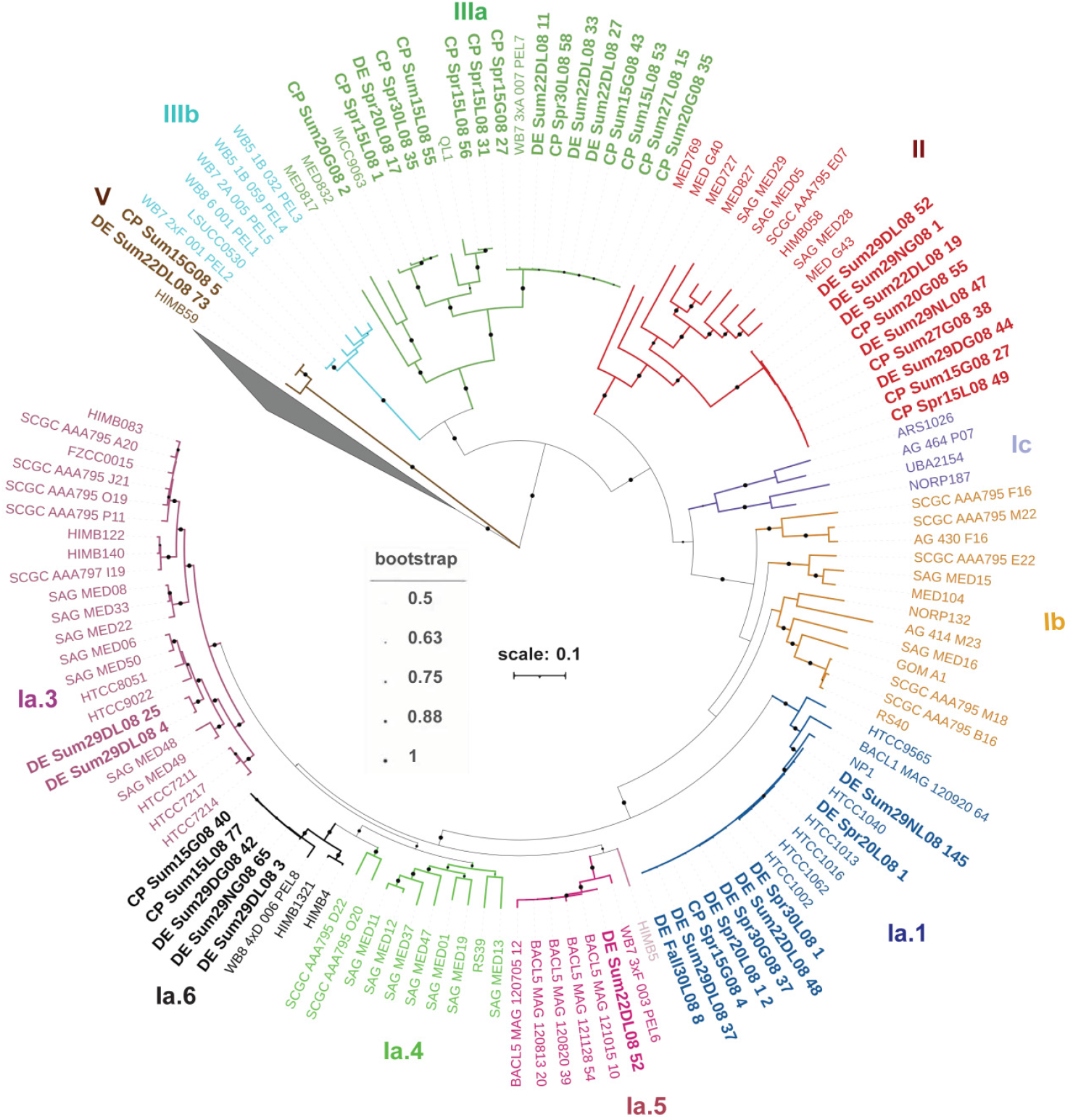
Phylogenomic analysis of SAR11 MAGs from the Delaware Bay (DE) and Chesapeake Bay (CP) compared to previously characterized SAR11 and Rickettsia sp. (grey wedge) genomes via the Maximum Likelihood method and JTT matrix-based model (79). A total of 38 shared genes were detected via Anvi’o (76) with amino acid sequences derived from 135 genomes, SAGs or MAGs. There were 6532 positions in the final dataset. The fraction of trees out of 100 in which the associated taxa clustered together are indicated. The tree is drawn to scale, with branch lengths measured in the number of substitutions per site. Information/accession number for each SAR11 genome is listed in **Table S1**.

### SAR11 abundance in the Chesapeake and Delaware bays

To estimate the relative abundance of SAR11 genomospecies in our samples, we mapped each MG and metatranscriptome (MT) to SAR11 MAGs and genomes dereplicated at a threshold of 95% average nucleotide identity (ANI). We then compared their predicted abundances to Kaiju-based estimates of total *Pelagibacterales* or bacteria in their corresponding MG and MT. *Pelagibacterales* dominated the microbial communities from mid- and high-salinity < 0.8 μm samples (21-61% abundance) and were less abundant in the > 0.8 μm samples (8-25% abundance) (**Fig. 2a, see Fig. S2 at 10.6084/m9.figshare.19701460**). The relative abundance of the genomospecies varied extensively and is detailed in the supplemental text. Briefly, SAR11 genomospecies Ia.1 and IIIa.2 were most abundant in the spring, genomospecies Ia.1 and IIIa.4 dominated Chesapeake spring samples, while genomospecies II, Ia.5, Ia.6, IIIa.4 and V dominated in summer in both bays (**Fig. 2b**). The average ratio of Pelagibacterales reads from recovered MAGs to total Pelagibacterales reads in their corresponding metagenomes was 75% and ranged from 60-95% in non-freshwater samples (**Fig. 2c**).

**Figure 2.**
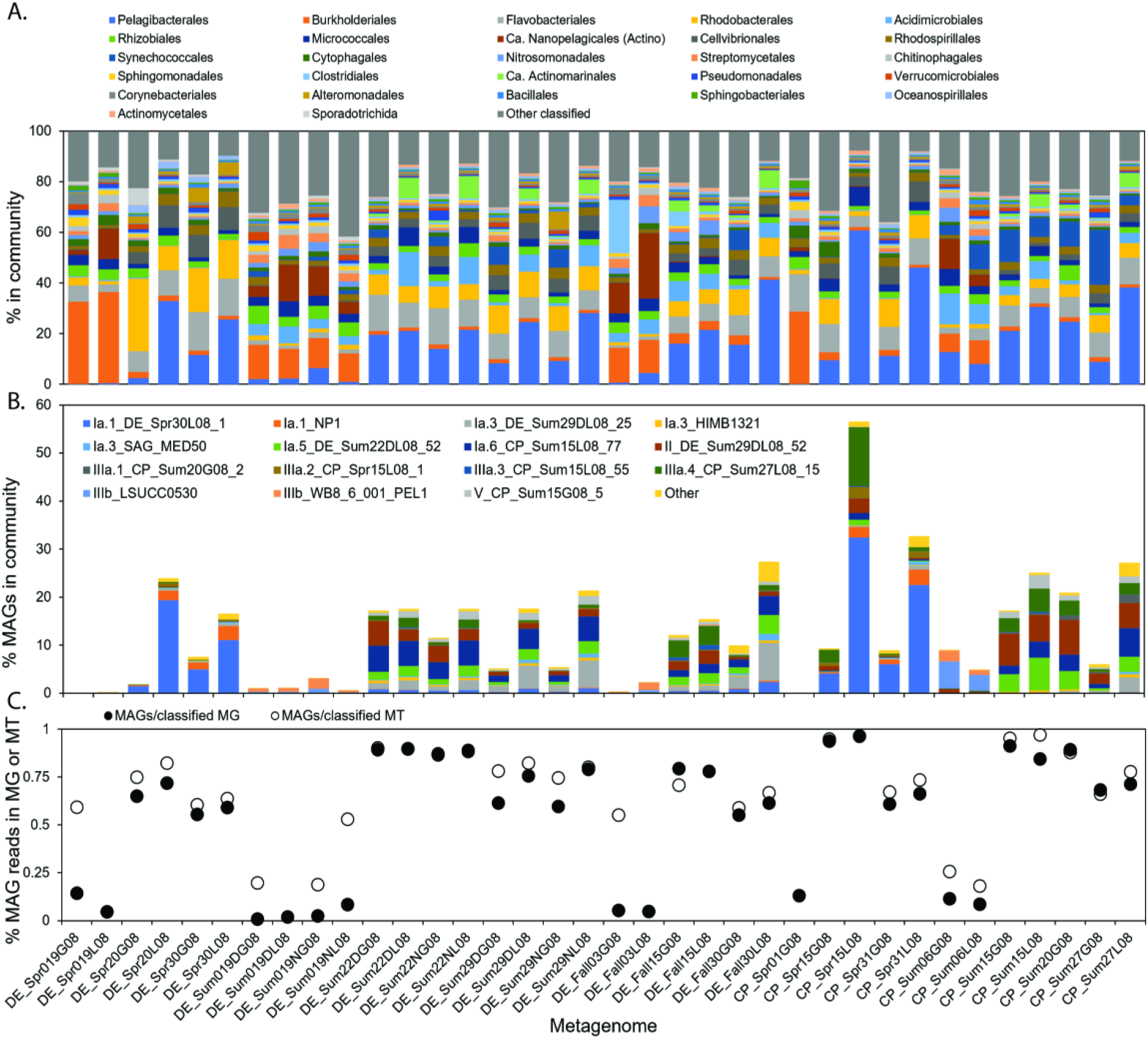
Percentages of Pelagibacterales in total classified Bacteria and Archaea in the DE and CP Bay metagenomes. A. Order level classification of metagenomic reads via Kajiu (87) in relation to total bacterial and archaeal reads. B. Percentages of representative Pelagibacterales MAGs/SAGs/genomes in the indicated metagenome in relation to total bacterial and archaeal reads. Abbreviations: CP= Chesapeake Bay, DE = Delaware Bay, Spr = Spring, Sum = Summer, the numbers following indicate salinity in PSU, G08 = >0.8 μm size fraction, L08 = <0.08 μm size fraction.

### Pangenome characteristics

Since SAR11 genomes/MAGs from these estuaries have not been comprehensively compared, we used Anvi’o to perform pangenome analyses of gene clusters, annotated KO (KEGG) genes, and COG genes. From 104 SAR11 reference genomes and medium to high-quality MAGs (>70% completeness and <5% contamination) (**Table S1**), 13,457 gene clusters were identified (see **SI at 10.6084/m9.figshare.19701478**). No gene cluster was shared between all 104 genomes, likely due to the incompleteness of the MAGs. However, 107 gene clusters, 254 KEGG Orthology (KO) terms and 349 clusters of orthologous genes (COGs) present in at least 50% of the genomes within a genomospecies were shared between genomospecies.

Since transporters make up a large percentage of SAR11 genomes, we next assessed their differences between subclades. We identified 97 transporter genes based on Pfam annotations (**Fig. S3**, see **SI at 10.6084/m9.figshare.19701478**). Transporter genes classified into 18 COG categories and their abundance varied by subclade. The number of genes in the G (carbohydrate transport and metabolism) COG category in each subclade varied in the order of highest to lowest number: V, Ia.3, II, 1a.5/Ia.6, Ia.1, respectively. The abundance of the ABC-type proline/glycine betaine transport system (*pro*VWX) was most represented in subclades Ia.3, Ia.5 and V (**Fig. 3**). Both ABC and TRAP transporters for carbohydrates were well represented in most subclades (**Fig. 3**, see **SI at 10.6084/m9.figshare.19701478**), although there were some notable differences. Most subclades had variable numbers of genes in the ABC-type glycerol-3-phosphate (*ugp*ABE) and ABC-type sugar transport systems (e.g., *malK*). The supplemental text discusses other transporter genes in more detail.

**Figure 3.**
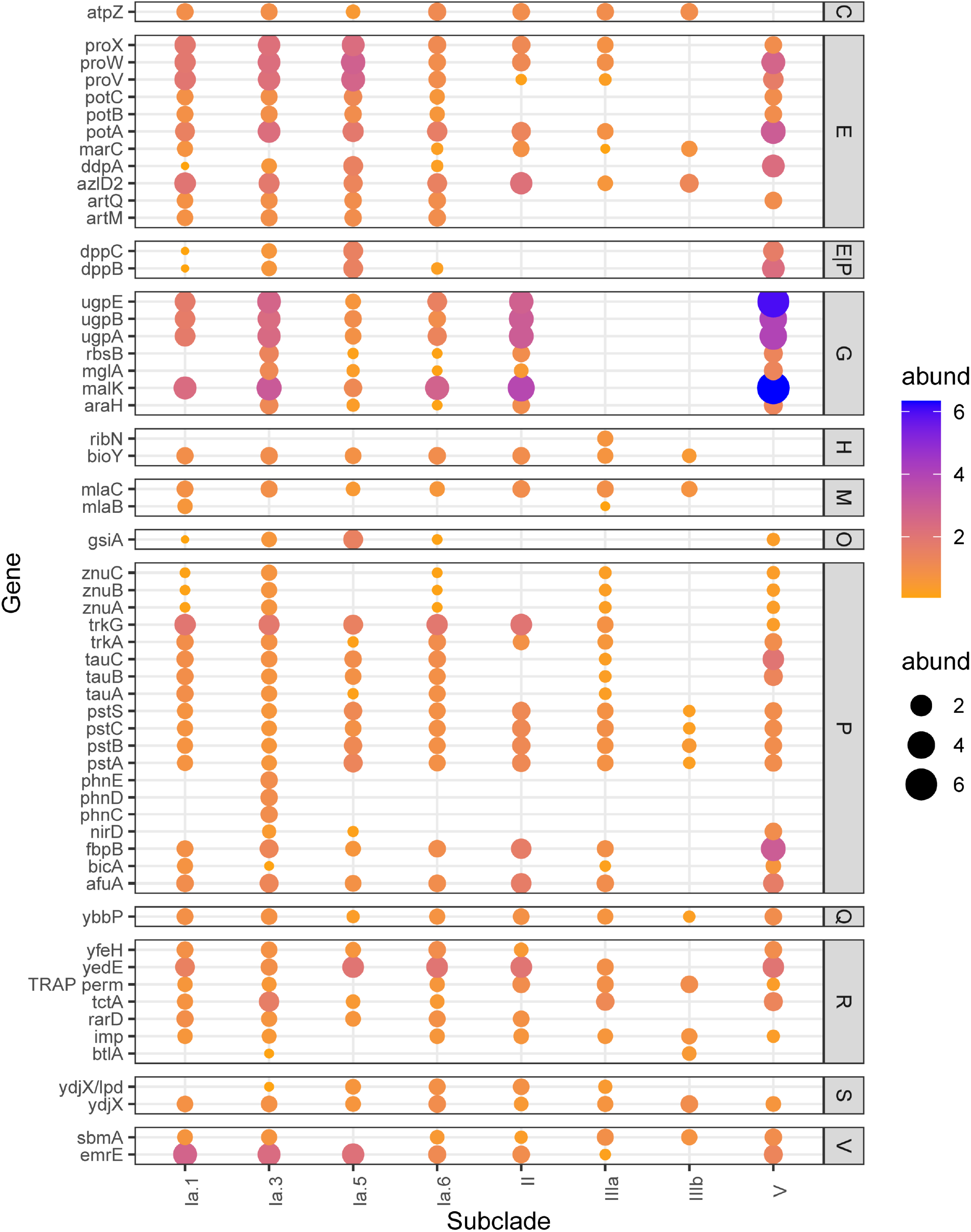
Average abundance of the indicated transporter gene in each SAR11 MAG. The transporters per genomospecies was filtered and sorted by the COG category indicated on the right. Only transporters found in 1-7 genomospecies are plotted. For a list of names and abundances of all transporters see **SI at 10.6084/m9.figshare.19701478**.

As with transporter genes, carbohydrate metabolism genes were differentially present between subclades and genomospecies. These included genes in the classical Embden-Meyerhof-Parnas (EMP) glycolysis pathway, Entner–Doudoroff (ED), a variant of ED pathway, Pentose Phosphate Pathway (PPP), tricarboxylic acid (TCA) cycle, and the glyoxylate shunt, which are known to differ between representative SAR11 genomospecies (13, 28). The two V and all II MAGs from the Delaware and Chesapeake, as well as the two IIIb genomes, contained all but one of the genes in the EMP pathway. None contained genes for the full ED pathway. However, MAGS from genomospecies Ia.5 and II and NP1 (Ia.1) contained all but one of the expected genes in the variant ED described in *Pelagibacter ubique* HTCC1062, and most genes were on the same contig (**Fig. 4, see SI at 10.6084/m9.figshare.19701478**). Ia.1 MAGs from the Delaware and Chesapeake were missing at least three genes previously described in the variant of the ED pathway, and the gene content and synteny was identical to that of HTCC1016. Members of the 1a.3, 1a.6, and IIIa subclades were missing 4 of 11 genes in this pathway (**see SI at 10.6084/m9.figshare.19701478**). Only MAGs and genomes within subclades IIIa, IIIb and V contained all the genes in the parallel glycolysis PPP. We next explored the genetic potential of SAR11 genomospecies for utilizing the TCA cycle or the alternate glyoxylate shunt. Almost half of the MAGs in the Ia.I subclade had identical glyoxylate shunt genes present in reference Ia.1 genomes. The two V MAGs contained most or all genes in the TCA cycle but not the glyoxylate shunt. Most representatives in IIIa and IIIb contained most or all genes in the TCA cycle and glyoxylate shunt. However, MAGs within II and Ia.3 genomospecies from the Delaware and Chesapeake bays were missing genes in glycolysis, the TCA cycle and the glyoxylate shunt, possibly due to incomplete genomes. Additionally, MAGs in genomospecies II, IIIa.2 and IIIa.4 contained, phasin and *phaC* genes, involved in carbon storage via polyhydroxyalkanoate (PHA) metabolism, used when there is an imbalance between carbon and nutrients in several bacteria (42). Across genomospecies, we also observed a clear difference between the presence of genes in many known C1 metabolism pathways in SAR11 (13, 22). The Ia.1 and V subclades contained most genes, except for carbon monoxide oxidation genes (*cut*MLS-*cox*G), which were present in some Ia.3, Ia.4, Ia.5, Ia.6, Ib, Ic, and IIIa subclades (**Fig. S4, see SI at 10.6084/m9.figshare.19701478**). Further details of the SAR11 pangenome is discussed in the supplemental text.

**Figure 4.**
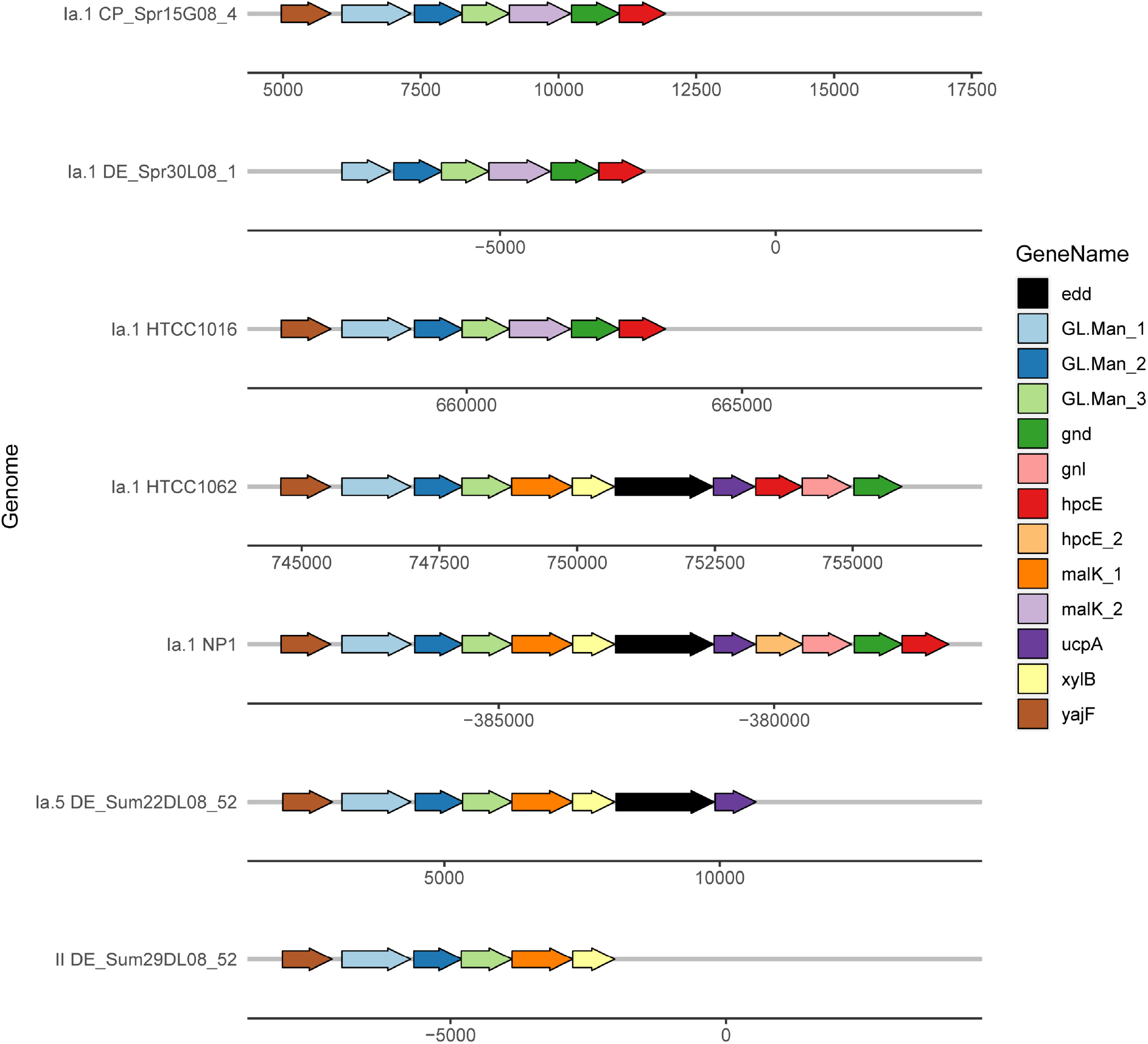
Gene neighborhood map of genes related to the alternate ED glycolysis pathway described in (28). Genomospecies and MAG or genome name listed on the left. Ia.5 and II MAGs also had genes on other contigs not listed here. Full names of genes corresponding to gene symbols are listed in **SI at 10.6084/m9.figshare.19701478**.

### Growth estimates

We used two independent methods to gain an understanding of the microbial growth rates across the representative MAGs/genomes of the genomospecies in our dataset. First, estimates of maximal growth rates for representatives based on codon usage patterns were below the minimum values in gRodon of doubling times > 5 hours, a known bias for slow growing organisms (43). Second, patterns of sequence coverage across the representative MAGs as assessed by CoPTR Peak to Trough ratios that reflect actual growth rates (44), varied between samples and representative genomospecies (**Table S2**). To broadly look at the entire SAR11 community in relation to transcript to genome ratios and nutrient concentrations, we first averaged CoPTR values in a sample. Average CoPTR values of all genomospecies within each metagenomic sample were correlated (r = 0.93, n = 11, p < 0.001) with the independently derived ratios of the percentage of total *Pelagibacterales* in the metatranscriptome in relation to the metagenome (MT/MG) from each sample (**Fig. 2c**). Additionally, both average CoPTR values and MT/MG were negatively or positively correlated with chlorophyll a, phosphate and ammonium in the spring and summer, respectively (**Figs. 5a-b, S5-S6**). Average CoPTR values were negatively correlated with nitrate in the spring, but not in the summer. We next evaluated the relationship between all CoPTR values and environmental variables (**Table S2, Figs. 5c-f, S7a**). All CoPTR values were significantly correlated with bay (Delaware>Chesapeake) and season (fall+summer>spring) (p<0.05, Wilcoxon rank), but were not with salinity. SAR11 genomospecies clustered into two separate groups based on their CoPTR values (**Fig. 6**). Group 1, with low, relatively constant values, included representatives from 1a.1, IIIa.2 and IIIa.4, while group 2, with higher and more variable values, included representatives from Ia.3, Ia.5, Ia.6, II and V. Average CoPTR values from multiple representatives within a genomospecies did not vary significantly (**Fig. S7b**). When the representative MAG CoPTR values were split into the two groups described above, none were significantly correlated with any Chesapeake environmental variable and only the second group, with representatives from genomospecies Ia.3, Ia.5, Ia.6, II and V, had values that were either positively (phosphate and silicate) or negatively (salinity) correlated with variables from the Delaware (**Fig. 5, S7**).

**Figure 5.**
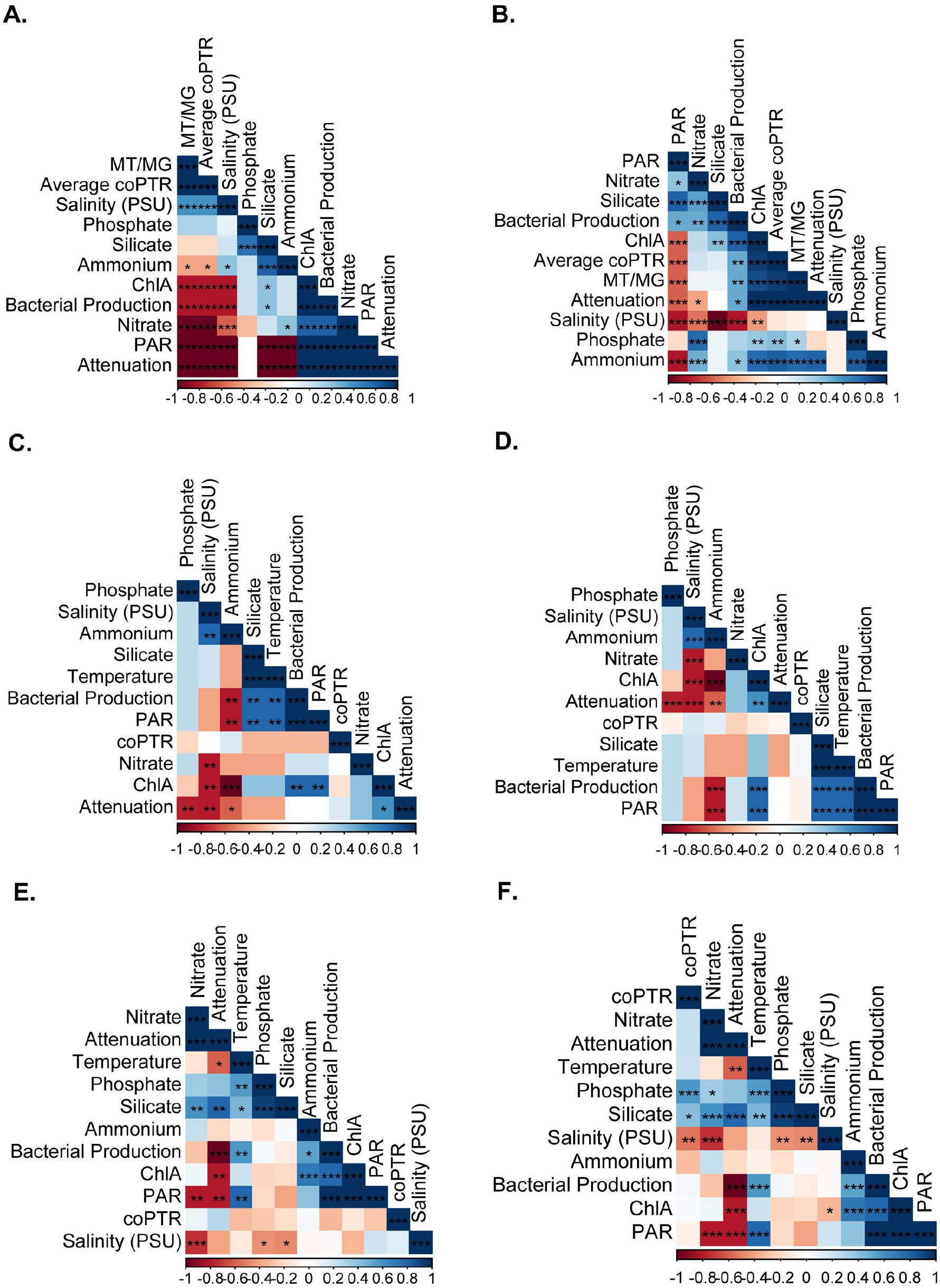
Correlations between environmental variables, including average CoPTR values and Pelagibacterales MT/MG ratios (A+B), and CoPTR values (C-F). A. Spring; B. Summer; C. Chesapeake, genomospecies Ia.1+IIIa.2+IIIa.4; D. Chesapeake, genomospecies Ia.3+Ia.5+Ia.6+II+V; E. Delaware, genomospecies Ia.1+IIIa.2+IIIa.4; F. Delaware, genomospecies Ia.3+Ia.5+Ia.6+II+V. * = p<0.05, ** = p<0.01, *** = p<0.001.

**Figure 6.**
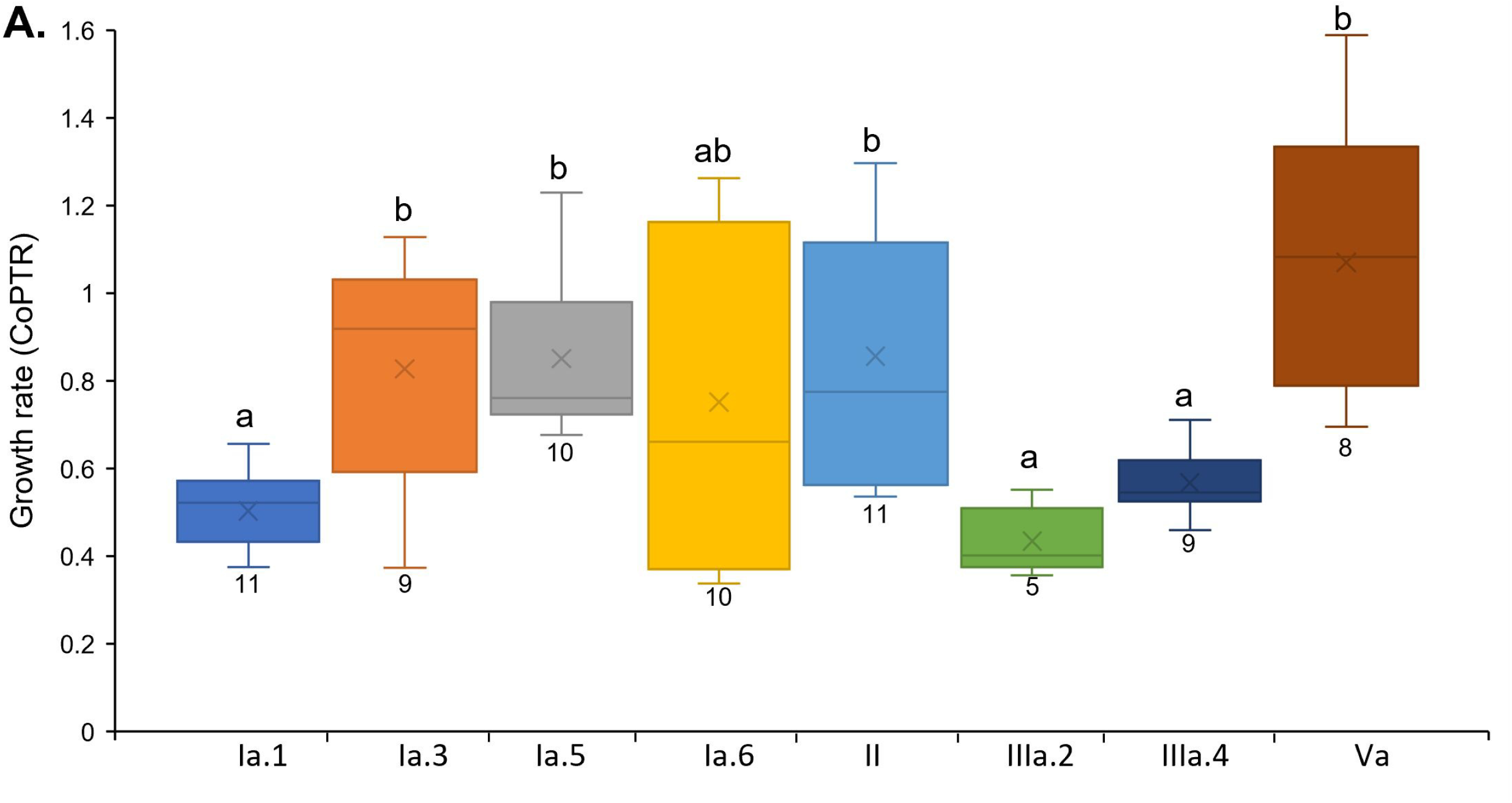
Peak to trough ratios of representative MAGs from the indicated SAR11 genomospecies in medium and high salinity L08 metagenome samples. Box and whisker plots of CoPTR values from the indicated genomospecies. The number of metagenomes is indicated below each box and the a or b or ab (not different) values above the boxes indicate significantly different groupings identified via the Wilcoxon sum rank test (95).

### Activity estimates

We first examined activity by looking at the relationship between the relative abundances of representative MAGs in MTs and MGs (**Fig. 7, see Fig. S8 at 10.6084/m9.figshare.19701463**). Representatives from genomospecies Ia.1 (NP1 only), Ia.3, Ia.5, and V had MT/MG ratios significantly greater than one, suggesting that they were more active than expected while the opposite was the case for IIIa.4 (p<0.02, Wilcon-signed rank test). The ratios of the other genomospecies were not significantly different.

**Figure 7.**
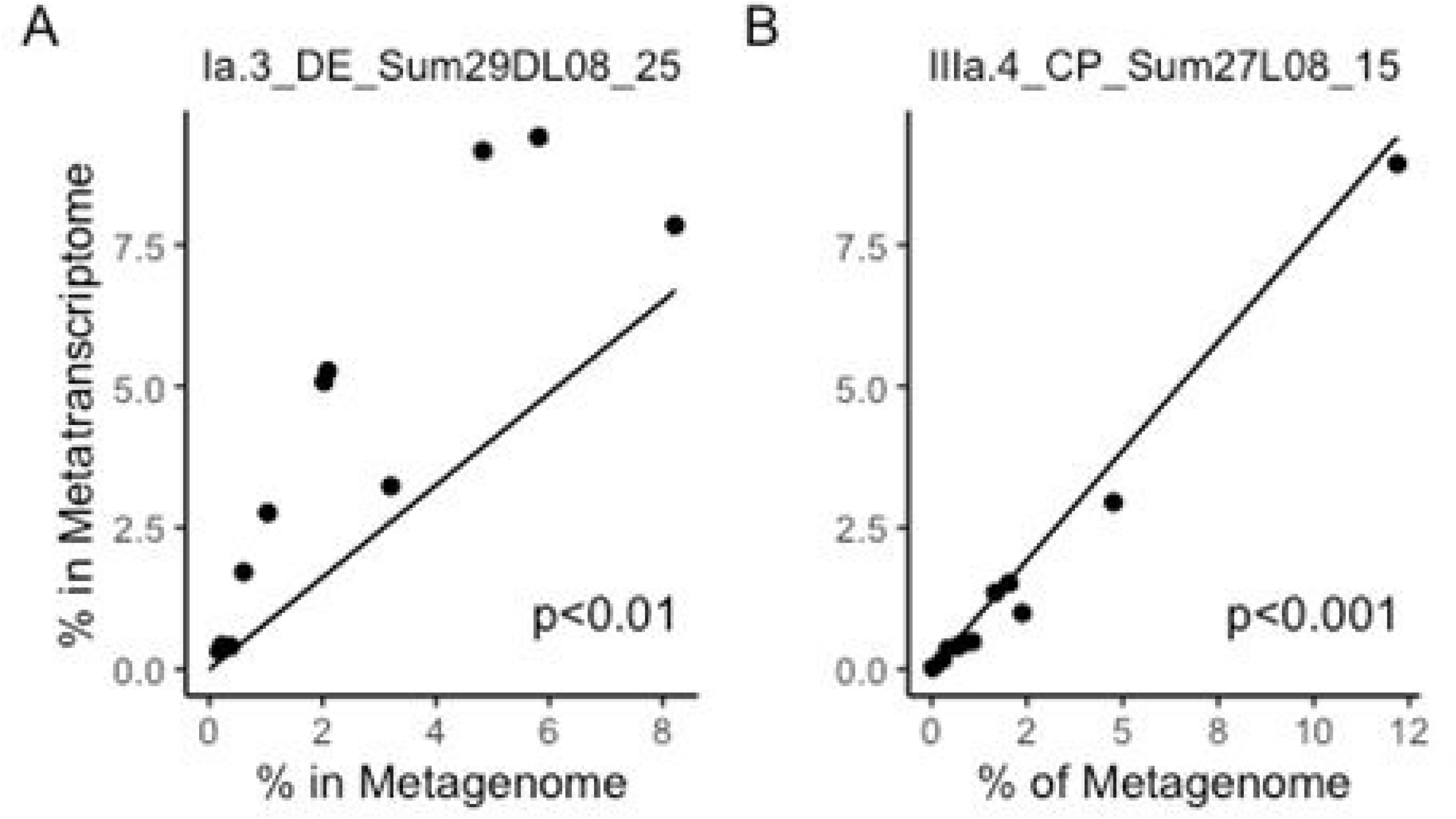
Relative abundance of two SAR11 genomospecies in metatranscriptome and metagenomic libraries. A. Genomospecies from Ia.3 subclade; B. Genomospecies from IIIa subclade. The p values are from the Wilcoxon signed-rank test of the significance that the ratio of MT to MG is not equal to one. See **Fig. S8 at 10.6084/m9.figshare.19701463** for data on other genomospecies.

Next, differences in transcription between conditions were evaluated from MT sequences mapped to gene clusters of representative MAGs listed in **Table S1.** Most of the observed variation was in the first axis (92%), which separated spring from summer samples (**Fig. 8**). The rest of the variation was related to bay and then salinity, especially with the Chesapeake MTs. We next compared transcript profiles from the representative MAGs between various conditions to examine whether there were commonalities in gene expression changes. The total percentage of genes from each MAG that either increased or decreased in each comparison ranged from 3.6-25.3% and averaged 11.6% (**see Table S3 at 10.6084/m9.figshare.19701457**). Gene expression differences due to salinity, especially in the Chesapeake, are detailed in the supplemental text (see **Fig. S9 at 10.6084/m9.figshare.19701469**).

**Figure 8.**
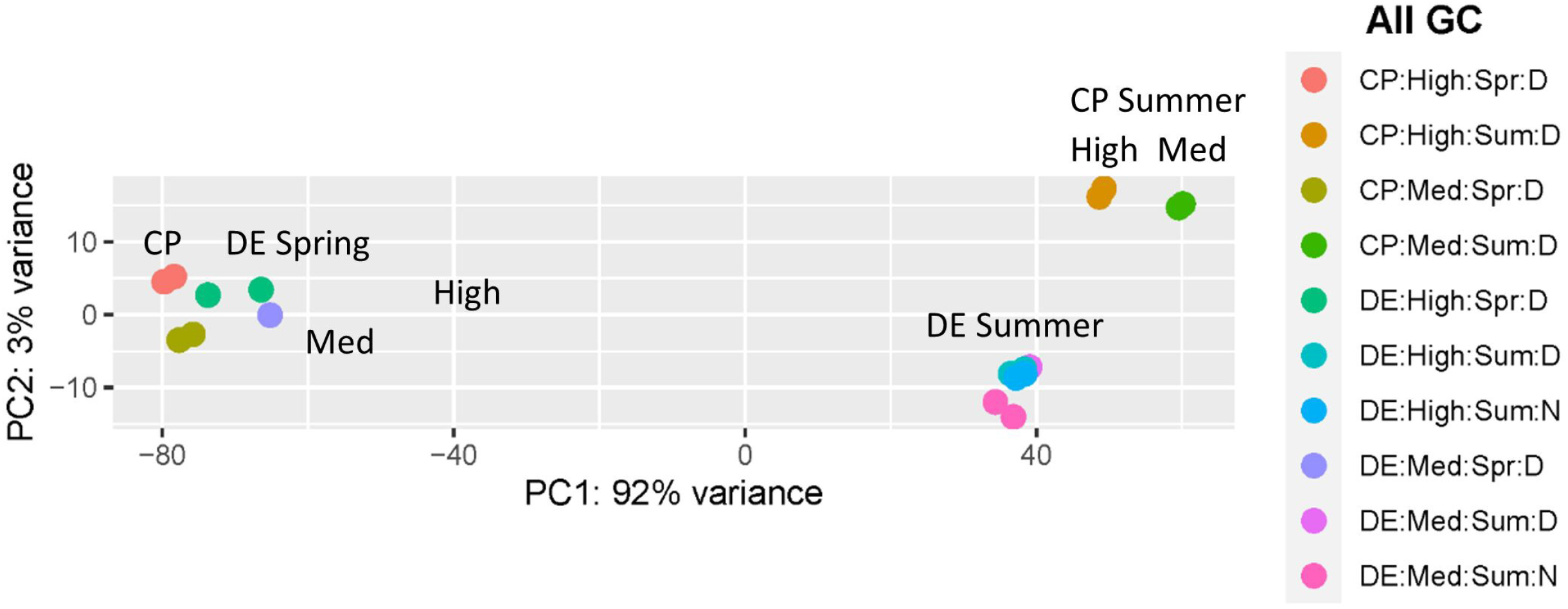
Principle Coordinate Plot comparing transcripts from the indicated samples at the gene cluster (GC) level. Representative SAR11 MAGs used are listed in Table S1. Transcripts were matched to gene clusters identified in Anvi’o (76) and normalized in DESeq2 (91).

Because we saw the most difference in growth rates of MAGs between the two bays, we concentrated on activity differences between the Chesapeake and Delaware, controlling for salinity and season. We compared the transcript counts mapped to representatives from genomospecies within subclades Ia.3, Ia.5, Ia.6, II, IIIa.4 and V between the Delaware and Chesapeake in the summer, separating medium salinity samples from high salinity samples to control for salinity (**Fig. 9, see SI at 10.6084/m9.figshare.19701484**). Expression of most genes in COG categories C (energy production and conversion), J (translation), O (post-translational modification, protein turnover, chaperone functions) and S (function unknown), including ribosomal protein genes, energy conservation genes and co-chaperonin genes, was higher in the Chesapeake than in the Delaware. In contrast, genes within the COG categories for amino acid transport and metabolism (E) and secondary metabolites biosynthesis, transport, and catabolism (Q) were generally upregulated in the Delaware compared to the Chesapeake. In general, expression of phosphate transport and regulatory protein encoding genes (*pstSCBA, phoBU*) and the ammonia channel protein encoding gene *amtB* was higher in the Chesapeake than in the Delaware, except for the phosphate transporters in genomospecies Ia.5, where the opposite was true. Expression of other genes previously associated with nitrogen limitation (45) besides *amt*B, namely *tauA* (taurine transport), *yhdW* (amino acid transport), and *glnA*, were also increased in the Chesapeake compared to the Delaware in the Ia.3 representative MAG (**see SI at 10.6084/m9.figshare.19701484**). While ammonium concentrations were much less in the Chesapeake compared to the high salinity waters in the Delaware, there was not much difference in the concentrations from the medium salinity samples (**Fig. S5**). Representative MAGs from genomospecies II and IIIa.4 had higher *phaC* and phasin gene expression in Chesapeake than in the Delaware. No differentially expressed genes in the glycolysis or the pentose phosphate pathway were observed from representative MAGs between the bays. However, expression of several TCA cycle genes was higher in the Chesapeake than in the Delaware in the V representative. Similar gene expression patterns in the Delaware summer L08 samples were observed when comparing high salinity and medium day salinity samples to the medium night salinity samples. These samples also had differences in growth rates and phosphate levels and are detailed in the supplemental text (**Fig. S10**).

**Figure 9.**
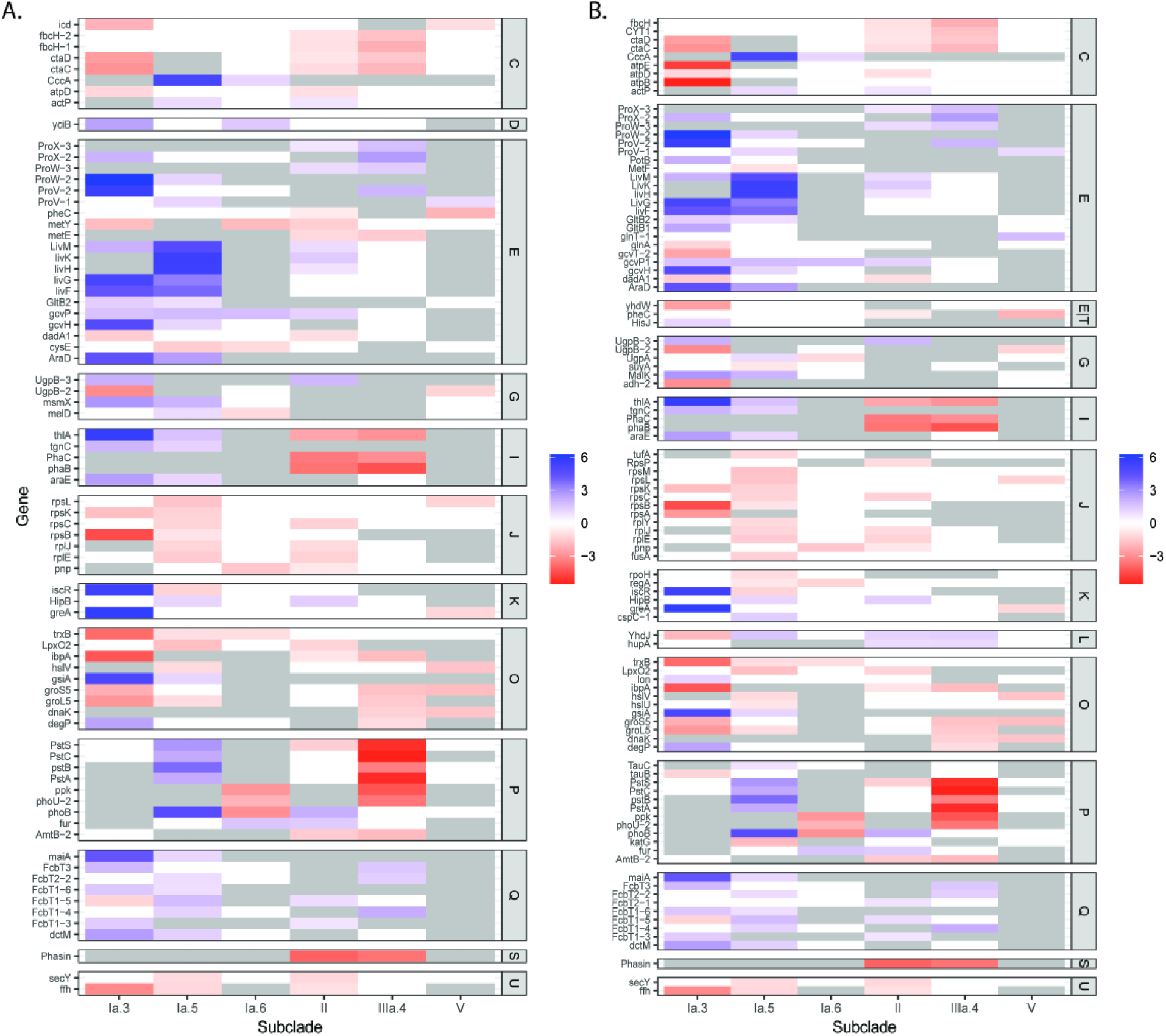
Common differentially expressed genes of representative MAGs from the indicated SAR11 genomospecies between Bay summer < 0.8 μm fraction (L08) metatranscriptome samples grouped by COG category. Log2 Fold Change values listed as determined by DESeq2 (91). A. High salinity, B. Medium salinity. Blue: increased in Delaware samples; red: increased in Chesapeake samples; white: not significantly different; grey: not present. COG categories are listed on the right. Gene names are defined in **SI at 10.6084/m9.figshare.19701478**.

## Discussion

Few estimates of growth rates of taxonomic groups below order level directly in natural environments exist beyond a few studies using MAR-FISH or rRNA:rRNA gene ratios (6, 25, 46, 47). Here, we found that growth rates of *Pelagibacterales* varied between and within genomospecies depending on environmental conditions, seasons, and bays. No other study has estimated SAR11 genomospecies *in situ* growth rates and compared them to environmental factors. Importantly, the average growth rate of genomospecies that dominated in the spring was negatively correlated with productivity measures and many environmental factors as one would expect with bacteria that dominate oligotrophic environments. However, the average growth rate of genomospecies that dominated in the summer was positively correlated with those same measures indicating different controls of growth rates. Phosphate concentrations and temperature, as well as transcripts involved in the cycling of phosphate and nitrogen compounds, were also positively and negatively correlated, respectively, to growth rates within the genomospecies that dominated summer samples. In addition to nutrients, variations in carbohydrate gene repertoires also helped explain differences in growth rates between these genomospecies. Finally, some genomospecies may be more suited to estuaries than open ocean systems. For example, at least one genomospecies in subclade II was not previously described and others in Ia.6 and IIIa not well represented in previous SAR11 ecogenomics studies done in oceanic waters and were found in estuaries (7, 48–50).

Past studies demonstrated generally low ratios of SAR11 rRNA to rRNA genes in coastal and estuarine environments (6, 26, 51–53), with some potential variation in estimated growth rates between seasons or salinities (6, 26). Others have developed algorithms using metagenomics or genomics to measure maximum potential growth rates, based on codon usage, as well as estimates of actual growth rates, based on differences between read depth at the origin of replication versus the terminus (PTR) (43, 44, 54). These would be ideal to use in environmental settings because they require input MAGs with high completeness and low contamination. However, both maximal potential growth rates and some measures of PTR are not sensitive, especially with slow growing organisms like those in the SAR11 clade (43). We assembled SAR11 MAGs with high completeness and low contamination using many metagenomic samples for use in our growth rate estimates (41). Our average estimates of growth rate for SAR11 Ia.1 MAGs was about 0.5 per day using CoPTR (44), similar to estimates of *P. ubique* in culture (~0.4 per day) (25, 55) and of the total clade in situ, which ranged from 0.1-1.8 per day (31, 56). Additionally, we found that our average PTR estimates per sample also correlated well with average ratios of mRNA transcripts per genome reads. The results parallel what was found with total RNA:DNA or rRNA:rRNA genes in many previous culture-based studies (57, 58), but were different from a previous study with *P. ubique* (25).

SAR11 is typically known as an oligotroph that thrives in low nutrient conditions. SAR11 subclade abundances vary under different environmental conditions (20, 21, 59). Our results from the spring indicate SAR11 has an oligotrophic lifestyle in both estuaries, where the average growth rates and transcripts per genome reads both negatively correlated with chlorophyll a and nutrient concentrations. As expected from previous studies in temperate coastal waters, slow-growing members of genomospecies in the Ia.1 subclade dominated both the total microbial community and the *Pelagibacterales* in spring (18, 20). However, we found the opposite was true in the summer, where average SAR11 growth rates in the two estuaries were positively correlated with chlorophyll a, phosphate, and ammonium, but negatively correlated with nitrate. The summer estuarine samples were dominated by members of Ia.3, Ia.5, Ia.6, II, IIIa.4 and V. Earlier studies indicated that some SAR11 subclades prefer environments with higher nutrients. For instance, subclade II members dominated mesopelagic zones at the Bermuda Time series station two months after mixing events, while many Ia.5 and IIIa populations dominate more eutrophic estuarine environments (6, 9, 18, 20, 49, 50). While the differences in genomospecies presence might be expected under different nutrient regimes, our results indicate that growth rates are also differentially affected by environmental conditions, at least in these estuaries.

We expected slower growth rates in the spring versus the summer based solely on temperature effects (60, 61). However, we did not expect SAR11 growth rates to be slower in the Chesapeake than in the Delaware. This difference could be due to the amounts and types of carbon available in the Delaware compared to the Chesapeake from late summer phytoplankton blooms of mixed eukaryotes, especially diatoms. Our data suggest eukaryotic phytoplankton blooms were more of a factor in the Delaware than the Chesapeake; chlorophyll a concentrations in the mid-salinity samples from the Delaware were two times higher than that in the Chesapeake while Cyanobacteria were much more abundant in the Chesapeake than in the Delaware. Additionally, carbohydrate transporter genes from representative SAR11 MAGs were differentially expressed between the two bays. These genes encoded proteins in the sn-glycerol 3-phosphate transport system, as well as other ABC and TRAP-type carbohydrate transporters. These results suggest that the types of carbon utilized by these different genomospecies varied with environmental conditions.

SAR11 genomospecies also differed in their carbohydrate transporter and metabolism genetic repertoires that were related to growth rate differences. The total number of carbohydrate transporter and metabolism genes, when corrected for genome size, was related to the average growth rate of the representative MAG. This suggests that SAR11 genomes with more access to carbohydrates can replicate faster. Furthermore, MAGs in genomospecies Ia.1 and IIIa had significantly lower growth rates than MAGs in genomospecies Ia.3, Ia.5, II and V. The MAGs in the former group lack genes in the typical EMP and ED glycolysis pathways and the alternate ED pathway found in coastal Ia.1 isolates (28), but had a gene content and synteny identical to HTCC1016, isolated from coastal Oregon (23). However, MAGs in the latter genomospecies have most or all the alternative ED pathway (Ia.3, Ia.5, II) or most genes in the EMP pathway (II and V). The lack of an alternative ED pathway in our estuarine/coastal Ia.1 MAGs is surprising, given the pathway was found in our NP1 Ia.1 representative. Previous studies indicated that this pathway was much more prevalent in coastal SAR11 than those found in the open ocean (28).

Growth rates of individual SAR11 genomospecies estimated from CoPTR and activities estimated from MT/MG ratios generally agreed with each other, especially for Ia.3, Ia.5, IIIa.4, and V, that were dominant in the summer samples. Ia.1 and IIIa.2, both abundant in the spring, had higher MT/MG ratios compared to CoPTR values. In contrast, representatives from Ia.6 and II dominant in the summer samples, had lower ratios than expected compared to CoPTR values. Differences in phage infection, defense mechanisms, or gene expression of highly abundant transcripts such as proteorhodopsin or different transporters may explain some of the unexpected differences (28, 33, 62, 63). Pelagiphages were abundant in the viral fraction from these samples, but there were no clear differences in abundance that corresponded to SAR11 abundance, nor were the phages identified to subclade level (64). Further characterization of the possible mechanisms leading to these differences may help explain this disconnect between predicted growth rates and transcript to genome ratios seen in this study.

The observed differences in growth rate variation within a SAR11 subclade are more difficult to discern. We observed a significant correlation between growth rate and a few environmental factors, namely phosphate, silicate, temperature (all positive) and salinity (negative), but only with representative MAGs from Ia.3, Ia.5, Ia.6, II and V subclades with higher growth rates. Phosphate concentration best explained PTR estimates with subclades Ia.3, Ia.5, II and V, and there was a negative relationship between expression of genes associated with phosphate uptake and PTR values. This indicated that phosphate, along with access to carbohydrates mentioned above, are likely key factors in regulating the variation in growth rates. Differences in phosphate and phosphonate transporter protein abundance have been observed between the oligotrophic Sargasso Sea and nutrient-rich upwelling areas (65, 66), but not on these finer temporal and spatial scales. A more recent study using incubations of microbial communities demonstrated that SAR11 increased growth rates with the addition of phosphate but not the elimination of grazers (29). Phosphate starvation or possibly increases in phosphonate also may lead to large changes in SAR11 metabolism. SAR11 growth under phosphate-limited but organophosphonate-replete conditions led to modifications of lipid content in response to phosphate starvation (24).

Besides phosphate and ammonium transporter gene expression having a negative relationship to growth rate estimates in subclades Ia.3, Ia.5, II and V, we did not observe any other genes that were consistently differentially expressed in relation to PTR values. Transcription of some nitrogen-associated genes (*glnA, gltB*) also generally increased with lower growth rates in our comparisons, but others previously found associated with nitrogen limitation in cultured SAR11 (*yhdW, occT* and *tauA*) were not consistently increased with low growth rates (45). Our results support the use of PHA storage mechanisms in two subclades (II and IIIa.4) under nutrient limiting conditions, where transcript levels of a PHA polymerase (*phaC*) were negatively correlated with growth rates in the representative MAGs. PhaC activity decreased during growth in *Pseudomonas putida* and in many bacteria increases under nutrient-limiting conditions (67–69). Transporters for some sugars and amino acids were increased in several MAGs with higher growth rates, This is in contrast to genes involved in translation, transcription, post-translational modification, protein turnover and chaperones and intracellular trafficking and secretion, which showed generally higher expression when growth rates were lower. However, several similar changes in global transcriptional patterns were observed between salinities in the Chesapeake, where no significant differences in growth rates were found nor any relationship between growth rates and environmental conditions. This suggests that some transcriptional responses may not be specific to growth rate.

Our results illustrate the different strategies that *Pelagibacterales* subclades utilize to successfully dominate bacterial and archaeal communities in a variety of estuarine and ocean systems. Not only do distinct members of the clade display niche specificity that is reflected in their genomic repertoire, but they also respond differently to environmental conditions as reflected by their growth rates. The differences were influenced by phosphate, ammonium, and carbohydrate transport and storage mechanisms. Our abilities to track these differences relied on a seasonal and salinity gradient study from two similar, but environmentally contrasting estuaries. This approach can be applied to understand the relationships in other estuarine microbial populations. Further research is needed in coastal or open ocean systems to examine SAR11 growth rates in relation to comprehensive space and time gradients.

## Methods

### Environmental Characterization and Sample Collection

Surface (~1.5 mbsf) water samples were collected using a rosette sampler with associated conductivity, temperature, depth (CTD) profiles during three cruises in 2014 and two cruises in 2015 of longitudinal transects of the Delaware and Chesapeake Bays, respectively, on the R/V Sharp (https://www.bco-dmo.org/dataset/565451). Water quality data was measured using a Sea-Bird data sonde, light intensity and attenuation were measured, and water samples were filtered for later measurements of nutrients and bacterial production as previously described (70–73). Collections for metagenomic and duplicate metatranscriptomic samples were identical to those previously described (41, 74).

### MAG taxonomic diversity and abundance

Surface water samples were collected from spring and summer salinity transects along the Delaware and Chesapeake Bays in 2014 and 2015, respectively (Supplemental text, **Fig. S5**) (41, 74). Metatranscriptome (MT) and metagenome (MG) sequencing, metagenome assembly, metagenome-assembled genome (MAG) generation, MAG quality assessment (completeness and contamination), and MAG taxonomic assignments are detailed elsewhere (41). Briefly, samples were collected, filtered to either 0.8 μm or through 0.8 μm and onto 0.2 μm size fractions, and total nucleic acids were extracted. Libraries were made from either DNA or RNA from single or replicate samples, respectively, at the Joint Genome Institute and sequenced. 44 MAGs classified as *Pelagibacteriales* with at least 80% completeness and less than 5% contamination determined by either CheckM (75) or Anvi’o (76) were used in this study (41) (**Table S1**). The MAGs were determined to be at least of medium quality due to the lack of rRNA genes in most of the genomes, according to the Minimum Information about a Metagenome-Assembled Genome standard (77). The phylogenetic position of these MAGs in relation to 88 NCBI reference genomes (78) was determined with Anvi’o phylogenomic analyses (76) with the Bacteria_71 dataset, where included genes needed to be present in at least 90% of the genomes. Subsequent phylogenomic maximum likelihood tree generation with 500 replicates was generated using the JTT model in MEGA X (79) and partial deletion from the resulting 38 ribosomal protein genes. The font sizes and colors were modified in iTOL from the resulting Newick tree (80). The average nucleotide identity (ANI) and average amino acid identity (AAI) between MAGs and reference genomes were calculated using MiGA version 0.7.26.2 using the --fast and --haai-p options (81). The AAI similarity matrix was plotted in heatmaply in R (82).

The above SAR11 MAGs and reference genomes were dereplicated at 95% identify by drep v2.6.2 and verified with MiGA version 0.7.26.2 (81, 83). Ribosomal RNAs were subtracted from MTs using SortMeRNA (84). Anvi’o v7 (76) MG and MT profiles were made by competitive mapping to representative dereplicated MAGs (**Table S1**) using Bowtie2 v2.3.4.1 (85) in default end-to-end mode (85). A count table of the resulting mapped reads and the Anvi’o gene-calls file reformatted to gff3 was generated using featureCounts v1.5.2 (86). In addition, MG and MT classifications to Bacteria-Archaea and order levels were performed with Kaiju v1.7.3 (87). The number of reads mapped to each group were used to estimate abundances and to correct read counts between samples.

### Pangenome analyses

Pangenomes were generated from MAGs and reference genomes/single amplified genomes /MAGs that were at least 70% complete and less than 5% redundant (**Table S1**) using a gene clustering approach (Diamond) in Anvi’o v7 (76). Protein-coding genes in each MAG were identified with Prodigal v2.6.3 (88) and annotated internally with KEGG, COG and Pfam profiles in Anvi’o v7 (76) and externally with Prokka v1.13 (89). Gene cluster, KEGG and COG counts for each MAG/reference genome were extracted from the anvi-summarize gene cluster table and re-formatted using the reshape2 package in R (90).

### Growth and Activity measures

Maximum growth rate were estimated with gRodon in R (43) using representative MAGs listed in **Table S1**. Rates for representative MAGs from each subclade from the fraction passing through a 0.8 μm filter (L08) were estimated via default CoPTR analyses (44), which estimates peak to trough ratios from mapped MG reads. Additional MAGs (**Table S1**) and *Pelagibacter ubique* (HTCC1062) were used in multiple replicate CoPTR analyses to validate the results.

The count table of MT reads mapped to representative MAGs described above was used for RNA-seq analysis in the R DESeq2 package (v3.4.3) (91). The count table was first condensed to gene cluster values (identified in Anvi’o) per MT sample, fitted with a negative binomial distribution and transformed by regularized log transformation in DESeq2. DESeq2 was also used to generate the principal component ordination plot. For differential expression analyses, subsets of the original count table were used for each representative MAG of a SAR11 subclade. Samples with more than ~50,000 reads mapped to the MAG were used in DESeq2 (91) to identify differentially expressed genes, which excluded particle-attached (cells captured on a 0.8 μm filter) samples because of the low abundance of hits. To further reduce any potential effects of abundance differences, comparisons were limited to within season, using SAR11 MAGs with less than two-fold change in relative abundance or coverage or less than one-fold or no significant differences in *gyrAB* gene expression (92, 93) compared to one another.

### Statistical analyses

To test for significant differences in CoPTR values between genomospecies as well as the MT to MG ratios that differed from 1, the Shapiro test (94) was first used to check for normality, followed by the non-parametric Wilcoxon rank sum test (95). All statistical tests were performed in R version 4.1.0 (96). Pairwise correlation matrices between CoPTR and environmental variables were calculated using the cor function in base R with Spearman as the model. The p-value for each pairwise correlation was calculated using the cor.mtest function in R’s corrplot library (97). The resulting correlation and p-value matrices were visualized using the corrplot function in the same R library. Linear regression analyses between CoPTR and environmental variables were performed using the ggscatter function in R’s ggpubr library (98). The non-parametric Spearman’s correlation coefficient was used in all correlation analyses.

## Acknowledgements

This work was supported by grants from the National Science Foundation (OCE-1261359) and the DOE (CSP-1621) to BJC and DLK and an NSF grant (EF-2025541) to BJC. We thank Matt Cottrell and Liying Yu for help with sampling and nutrient analyses and the crew of the R/V Hugh R. Sharp for sample collection. We also thank Tijana Galvina del Rio at the Joint Genome Institute for project help.

BJC and DLK obtained funding for the study. All authors helped with sampling. BJC prepared the samples and analyzed the data, with help from SJL. BJC wrote the paper with input from SJL and DLK.

## Supplemental Legend (for Supplemental Text, Tables S1 and S2, Figures S3, S4, S5, S6, S7 and S10, all others linked to figshare)

**Supplemental Text** – includes additional results and discussion ancillary to the main text.

**Table S1.** Properties of SAR11 MAGs, SAGs and genomes used in this study. Subclades were defined based on identities to previously characterized MAGs, SAGs and genomes via phylogenomics and AAI/ANI analyses. Genome/MAG properties were determined with Anvi’o.

**Table S2.** CoPTR values from a representative MAG in the indicated subclade from Delaware (DE) and Chesapeake (CP) Bay metagenomes from the less than 0.8 μm size fraction. The number after Spr (Spring), Sum (Summer or Fall is the salinity in PSU. N = night, D = day. Representative MAGs are listed in **Table S1.**

**Figure S3.** Number of transporters in the indicated COG category in each SAR11 subclade. Values are normalized to average genome size. A complete listing of all MAGs and reference genomes in each subclade is listed in **Table S1**.

**Figure S4.** Average abundance of the indicated gene associated with C1 oxidative metabolism in each SAR11 subclade. Heatmap was filtered and sorted by the C1 metabolic pathway indicated on the right. For a full definition of gene symbols and list of gene names see **SI at 10.6084/m9.figshare.19701478**.

**Supplemental Figure 5**. Environmental Data for metagenomes and metatranscriptomes used in this study. For further details, please see https://www.bco-dmo.org/dataset/565451. Average temperatures (°C) for each bay and season are listed in the panels with production and PAR numbers. There are no samples collected in the Fall in the Chesapeake Bay.

**Supplemental Figure 6.** Correlations between Pelagibacterales reads in MT/MG or average CoPTR values of representative MAGs from each subclade (**Table S1**) and the indicated environmental variable in spring or summer samples. p values were calculated by cor.mtest, after determining R values in cor via the Spearman method.

**Supplemental Figure 7.** Growth rate estimates via CoPTR values. A. Correlations between CoPTR values of representative MAGs from each subclade (**Table S1**) and the indicated environmental variable in spring, summer and fall samples. B. Average CoPTR values from replicate MAGs from each indicated subclade. p values were calculated by cor.mtest, after determining R values in cor via the Spearman method.

**Supplemental Figure 10.** Common differentially expressed genes of representative MAGs from the indicated SAR11 subclades between DE Bay summer less than 0.8 μm fraction (L08) metatranscriptome samples grouped by COG category. Log2 Fold Change values listed as determined by DESeq2 (91). Blue: increased in DE day medium and all high salinity samples; red: increased in DE medium night samples; white: not significantly different; grey: not present. A. COG categories: C, C|T, E, E|G, E|T, G, H, I, P, Q and S. B. COG categories: D, F, J, K, L, M and O. COG categories are listed on the right. Gene names are defined in **SI at 10.6084/m9.figshare.19701478**.

