## Supplemental Text for "Controls of SAR11 subclade abundance, diversity, and growth in two Mid-Atlantic estuaries"

Running Title: SAR11 in two Mid-Atlantic estuaries

Barbara J. Campbell^1#^, Shen Jean Lim^1*^, David L. Kirchman^2^

^1^Department of Biological Sciences, Clemson University, Clemson SC, 29634

^2^School of Marine Science and Policy, University of Delaware, Lewes, DE 19958

**Supplemental Results/Discussion**

*Environmental conditions*

Surface water samples were analyzed from spring and summer salinity transects along the Delaware and Chesapeake Bays in 2014 and 2015, respectively (**Fig. S5**). Data from April 2015 Chesapeake samples were reported previously (1). Water temperatures averaged about 5 °C cooler in the Delaware in March 2014 compared to the Chesapeake in April 2015, 4.3 ± 0.1 vs. 9.8 ± 1.2, respectively. The Chesapeake was slightly warmer (~27 °C) than the Delaware (~24.7 °C) in August 2015 vs. August 2014, respectively. Cell counts were between 2 and 10 times higher and bacterial production averaged 4 times higher in the summer compared to the spring from both bays. Nutrients varied between seasons and salinities and were generally lower in the Chesapeake compared to the Delaware. For instance, nitrate decreased from >170 to less than 1 μmol/L along the salinity gradient in summer Delaware samples, whereas it ranged from 20 μmol/L to less than 7 in the summer Chesapeake transect. Ammonium was less than 1 μmol/L in the spring and summer mid-salinity samples from the Chesapeake, but all other samples had concentrations greater than 3.7 μmol/L. Phosphate concentrations ranged from undetectable in spring mid- and high-salinity and summer mid-salinity Chesapeake samples to greater than 0.3 μmol/L in all low salinity samples. These results indicate possible phosphate limitation in spring, especially in the Chesapeake. In addition, lower nutrients in the Chesapeake compared to the Delaware summer samples supports previous observations of warm, low nutrient and stratified waters in the Chesapeake, while Delaware waters were mixed (2-5).

*MAG taxonomic diversity and abundance*

SAR11 Ia.1 included 9 MAGs from the Delaware and Chesapeake, *Pelagibacter ubique* HTCC1062, and other previously identified Ia.1 strains, which were further subdivided into at least 3 genomospecies. Ia.3 included two Delaware MAGs that are most related to members within the previously defined Ia.3/VIII subclade (6, 7). The third, Ia.4, did not contain any MAGs from the Chesapeake or Delaware. The fourth, Ia.5, includes two Delaware MAGs and several MAGs from the Baltic Sea and one from the Chattahoochee ecosystem (8, 9). The last Ia subclade, Ia.6, a new subgroup that contains five MAGs from the Delaware and Chesapeake, as well as a MAG from the Chattahoochee River and three marine isolate genomes previously classified within the Ia.3 genomospecies (6, 7, 9). All but one of those isolates, HIMB5, clustered in the Ia.6 subclade by hAAI (**see** **Fig. S1 at 10.6084/m9.figshare.19701379**). HIMB5 clustered with Ia.3 members. Estuarine *Pelagibacterales* Chesapeake and Delaware MAGs identified in subclade II were distantly related to other subclade II MAGs (**Fig. 1, see** **Fig. S1 at 10.6084/m9.figshare.19701379**). Chesapeake and Delaware MAGs within subclade IIIa were generally distantly related to one another, and three of the four genomospecies had representatives from both bays. There were two MAGs, one from each bay, within the V subclade.

Besides Pelagibacterales, other highly represented groups in the Delaware and Chesapeake Bay samples were *Burkholdariales*, *Actinobacteria*, *Flavobacteriales*, *Rhodobacterales* and *Synechococcalaes* (**Fig. 2a**). The lowest relative abundances of *Pelagibacterales* were in freshwater samples, ranging from 0-13% (**see Fig. S2a at 10.6084/m9.figshare.19701460, Fig. 2a**). Total dereplicated MAGs, including representative IIIb genomes not found in our filtered MAGs (**Table S1**), made up between 0 and ~58% of all classified *Pelagibacterales* (**Fig. 2b, see Fig. S2a at 10.6084/m9.figshare.19701460**). Representative MAGs and genomes from the SAR11 Ia.1 subclade dominated spring samples from both bays, with an abundance up to 32% in the Chesapeake Bay and were generally greater in mid- than high-salinity samples (**Fig. 2b**). The abundance of SAR11 Ia.1 was less than 2.3% in the summer and fall. Subclade IIIa.2, which comprised 0.01-2.4% of the community, was highest in the Chesapeake Bay spring samples. Genomospecies within subclades Ia.5, II, and V, which ranged in abundance from 0.02-6.7, 0.02-7.7 and 0-2.8% of the community, respectively, were most abundant in mid-salinity summer samples. Genomospecies within subclade IIIa.4 was most abundant in Chesapeake spring and summer mid-salinity samples (range 0-12.2). Genomospecies within Ia.6 and Ia.3 were highest in summer and fall high-salinity samples, where they peaked at 5.9% relative abundance.

To evaluate the abundance of MAGs to total *Pelagibacteriales*, we used Kaiju identified bacterial and archaeal sequences, and calculated the percentage of reads mapped to dereplicated *Pelagibacterales* MAGs to the total reads classified as *Pelagibacterales**,* which averaged 75% (**Fig. 2c**). The percent of recovered SAR11 MAGs ranged from 0.007-0.14% in freshwater samples, which only included previously published IIIb representatives (9, 10). The number of MG or MT reads classified as *Pelagibacterales* and those mapped to *Pelagibacterales* MAGs in relation to all Kaiju classified bacterial and archaeal sequences were generally similar (**see Fig. S2 at 10.6084/m9.figshare.19701460**). However, besides one outlier, Fall > 0.8 µm (G08), the ratios varied (range 0.39-1.69), with spring Delaware samples and all Chesapeake samples generally below 1, and with Delaware summer samples above 1.

Our data indicate that members of the SAR11 clade were very abundant and diverse in the Delaware and Chesapeake. Previously, we estimated relative abundances of up to 60% in a summer transect of the Delaware by 16S rRNA gene amplicon analysis (11, 12). Here, only the spring samples from the Chesapeake contained between 45-60% of *Pelagibacterales*. The abundance of this clade in all other samples averaged about 25% in mid and high salinity samples and much less in freshwater samples, similar to other estimates in marine and freshwater ecosystems (9, 13, 14). The abundance of this clade in the Delaware and Chesapeake estuaries is surprising, based on the generally eutrophic nature of estuaries and what others have observed in other estuaries (2, 8, 11, 14, 15).

Based on our and other previous estuarine studies (8, 9, 11, 12, 14, 16), we were not surprised to discover representatives from SAR11 subclades Ia.1, Ia.3, Ia.5, Ia.6, IIIa and IIIb in the Delaware and Chesapeake estuaries. However, the extensive diversity of subclade IIIa was somewhat unexpected. Genomospecies IIIa.2, mostly closely related to IMCC9063 (17) was only found in the spring, while IIIa.4, related to a PEL7 MAG in the Lake Lanier estuarine system (9), was abundant in the spring and summer mainly in the mid-salinity samples. Genomospecies IIIa.1, related to marine representatives of IIIa (18), was mostly found in higher salinity samples, but in low abundance. We also found seasonal patterns in representatives from Ia.3, Ia.5, Ia.6 and V, which were all abundant mainly in the summer. New to this study was the presence of a novel genomospecies within subclade II, which made up to 7% of the total bacterial and archaeal community in the summer. Subclade II genomes were previously represented by SAGs generated from deeper marine waters from the oxygen minimum zone of the ETNP and surface waters from the Gulf of Mexico or MAGs from the TARA oceans sampling of the Mediterranean Sea, (7) and have not been recognized in estuaries previously (9).

*Pangenome analysis*

Transporters were very abundant in the MAGs, with anywhere from about 60-125 per 1.3 Mb genome size (**Fig. S3**). Most of the transporters classified by COG were of the ABC type and were shared between all MAGs (**see** **SI at 10.6084/m9.figshare.19701478**). There were several TRAP-type transporters as well. Several notable differences in some transporter genes between the genomospecies were observed (**Fig. 3, see** **SI at 10.6084/m9.figshare.19701478**). The peptide/nickel transport system (*dpp*BC) was present only in Ia.3, Ia.5 and V. While all subclades contained ammonium (amt) and inorganic phosphate transporters (*pst*ABCS), only representatives in Ia.3 contained transporters for organic phosphate, or phosphonate (*phn*CDE). The ABC-type Mn2+/Zn2+ transport system (*znu*ABC) was absent from Ia.1, Ia.5, II, and IIIb SAR11 subclades, and mostly absent from Ia.6. The ABC-type taurine transport system (*tau*ABC) was also absent in subclade IIIb, as well as II, and weakly represented in IIIa. This variability in transport gene content among SAR11 subclades was expected (19, 20) and likely contributes to ecological niches observed in these estuaries and other marine environments (21-26).

More C1 metabolism genes were found in pangenomes from subclades Ia and V than those from subclades II or IIIa (**Fig. S4**). Carbon monoxide genes (*cut*LMS) were present in representative Delaware or Chesapeake MAGs (**Table S1**) from subclades Ia.5, Ia.6 and IIIa.4. Most of these C1 compounds, such as methanol and carbon monoxide, were likely derived from phytoplankton and other microbes and generally present in low concentrations in marine waters (27-30). Pangenomes for Ia.1, Ia.3, and V subclades contained most or all genes necessary for tetrahydrofolate-linked oxidation, methanol oxidation, and methylamine oxidation. Paralogous aminomethyltransferase genes, including *dmd*A, involved in dimethylsulfoniopropionate (DMSP) demethylation (31) were also detected in these pangenomes. Subclade Ia.1 representatives encoded a near-complete glycine-betaine (GBT) oxidation pathway (32), including *bhm*T, *dmg* and sarcosine dehydrogenase genes (**see** **SI at 10.6084/m9.figshare.19701478**). Up to three copies of the L-proline glycine betaine transport gene, *pro*X (33), were identified in all MAGs (**see** **SI at 10.6084/m9.figshare.19701478, Fig. S4**). We could not confirm the presence of the glutathione dependent pathway that converts formaldehyde to CO_2_ previously identified in HTCC 7211 (32). Finally, the Ia.3, Ia.5, Ia.6 and summer IIIa subclades from the Delaware and Chesapeake Bays contained carbon monoxide oxidation genes (*cut*MLS-*cox*G) previously reported within the same subclades (19). These were not identified in the clade II and spring IIIa subclades. We also found the same genes in the same order in MAGs in the Ia.4, Ib, and Ic subclades, further expanding the presence of this metabolism in the SAR11 clade. CO, a product of chromophoric DOM transformed by UV light (34), is present in the surface ocean at maximum concentrations in mid-day summer, but is also found in and produced from particulate organic matter, especially in the fall (35). SAR11 subclades that have this set of genes may gain energy via respiratory oxidation of CO to CO_2_ without biomass incorporation (36, 37). Our results suggest that CO oxidation may be an important contributor to the growth of some warm water SAR11 populations, as is hypothesized for other marine bacteria (36).

Glycine-serine auxotrophy, previously described in *P. ubique* (38), was not universally present in the subclades because genes necessary to make glycine and serine (*ser*ABC, glyoxylate precursor, threonine dehydrogenase, glycine hydroxymethyltransferase, *gcv*THP, *glc*B) were found in subclades IIIa and IIIb; subclade V was missing *ser*B (**see** **SI at 10.6084/m9.figshare.19701478**). However, threonine aldolase and 2-amino-3-ketobutyrate coenzyme A ligase were missing in IIIa, IIIb, and V.

*Differential gene expression analyses and growth rate patterns*

Similar expression patterns to the Delaware versus Chesapeake summer comparisons were observed when comparing the Delaware summer L08 samples from the high salinity and medium day salinity samples to the medium night salinity samples (**Fig. S10, Table S2, see SI at 10.6084/m9.figshare.19701484**). In this comparison, phosphate transporter gene expression was consistently increased in medium and all high salinity samples compared to medium night samples, and negatively associated with phosphate levels (**Fig. S5**). Expression of some genes involved in nitrogen compound uptake (*amt*B, gltB and *gln*A) was generally higher in parallel with phosphate transporter levels in Ia.3, Ia.5, Ia.6 and V (**Fig. S10, see SI at 10.6084/m9.figshare.19701484**). However, other transporters involved in nitrogen compound uptake, namely the transporters for amino acids, opines, and taurine, *yhd*W, *occ*T and *tau*A, were increased in several MAGs (Ia.3, Ia.5, Ia.6, IIIa.4) in the medium night samples compared to the medium day and all high salinity samples. Ammonium concentrations were about 3.5 times higher in the medium salinity night sample compared to the rest (**Fig. S5**). Additionally, there were no significant differences in expression of genes in glycolysis, the pentose phosphate pathway, the TCA cycle, including the glyoxylate shunt from different representative MAGs between these conditions (**see SI at 10.6084/m9.figshare.19701484**).

In addition to transcriptional comparisons with MAGs from the summer samples, we also looked for differences in transcription between salinities within the same bay and season to look for salinity effects on transcription as well as with the Delaware summer L08 samples from the high salinity and medium day salinity samples to the medium night salinity samples conditions as another comparison for growth rate differences. Expression differences between salinities were most apparent in the Chesapeake (See **Table S3 at 10.6084/m9.figshare.19701457, Fig. S9 at 10.6084/m9.figshare.19701469**). In fact, between 0-1.1% of genes were differentially expressed in different MAGs between salinities in Delaware samples, while between 2.9-25.4% of genes were differentially expressed in MAGs between salinities from the Chesapeake. Transcripts in COG categories C (Energy production and conversion), D (Cell cycle control and mitosis), F (Nucleotide metabolism and transport), J (Translation) and K (Transcription) were mainly increased in high salinity samples compared to medium salinity from MAGs present in the Chesapeake spring samples, except for some transcripts in NP.1, a member of subclade Ia.1 (**Fig. 1,** **see** **Fig. S9 at 10.6084/m9.figshare.19701469**). Expression of most genes involved in osmolyte regulation (*pro*XW) were increased in medium compared to high salinity samples, with the expression of *pro*XW-3 in genomospecies II, IIIa.2 and IIIa.4. Additionally, expression of *pha*C, and phasin, involved in PHA storage (39-41), were increased in IIIa.2 in medium compared to high salinity samples, while the opposite was true with *pha*C or phasin transcripts in IIIa.4 and II, respectively. Two of the most differentially expressed gene groups, ammonium (*amt*B) and phosphate transporters and regulators (*pst*BSAC, *pho*U), were increased in medium and high salinities, respectively. Here, phosphate concentrations were undetectable in both salinities. Ammonium levels were about 10 times higher in the high salinity compared to the low salinity samples (**Fig. S5**). Our results with *amt*B expression paralleled that in cultured cells, where increased *amt*B expression by cultured SAR11 was found in nitrogen-limited media (42).

While there was not much difference in CoPTR values between mid and high salinity spring Chesapeake samples with averages of 0.58 and 0.63, respectively, MAGs from Ia.1, Ia.5, Ia.6, II and IIIa.4 genomospecies all had slightly higher values in the high versus the mid salinity samples. We hypothesize that, based on estimated growth rates, transcriptional patterns and nutrient concentrations, ammonium and not phosphate was likely the limiting nutrient in the spring medium salinity samples, in addition to a probable imbalance of carbon compounds.

There were several transcriptional patterns from the MAGs present in the summer Chesapeake samples which were opposite that of the spring (**see** **Fig. S9 at 10.6084/m9.figshare.19701469, SI at 10.6084/m9.figshare.19701484**). For instance, transcripts in COG categories J (Translation) and K (Transcription) were mainly increased in medium compared to high salinity samples. Transcripts increased in medium compared to high salinities were the PHA storage gene, *pha*C, in genomospecies II and IIIa.4, and genes involved in amino acid metabolism, *leu*C (leucine metabolism), *gln*A (glutamine synthetase) and *gcv*T (glycine cleavage). However, transcripts in COG categories C (Energy production and conservation), Q (Secondary structure), V (Defense mechanisms), D (Cell cycle control and mitosis), F (Nucleotide metabolism and transport), L (replication and repair) and O (Post-translational modification, protein turnover, chaperone functions) were mainly increased in high compared to medium salinity samples. Proteorhodopsin gene expression was also increased in high compared to medium salinities from representatives in genomospecies Ia.5, II, and IIIa.4. Other expressed genes commonly increased in high compared to medium salinities were *kat*G (catalase), *amt*B (ammonium transport) and *glt*B (glutamate synthase), *sox*B (sarcosine oxidase), *dad*A (glycine cleavage), and *gln*T2 (glutamine synthetase), all involved in amino acid metabolism. Nitrate and ammonium concentrations were increased in medium compared to high salinity samples (**Fig. S5**). Branched chain amino acid transporters (*liv*FHM) were increased in Ia.5 in high compared to medium salinities, but the opposite pattern was observed in IIIa.4. As expected, the genes whose protein products are involved in osmolyte regulation [*pro*XW-3 (Ia.5, II, IIIa.4) and *pot*D (Ia.5 and Ia.6)] were increased in high compared to medium salinity samples. There were no significant differences in phosphate transporter gene expression between the salinities, although the differences in phosphate levels was high **(Figs. S5, see** **Fig. S9 at 10.6084/m9.figshare.19701469**). It could be possible that the phosphate concentration measured in the medium salinity sample was incorrect.

In contrast to the Chesapeake spring salinity comparisons, we did not observe consistent or significant differences in predicted growth rates in MAGs between medium and high salinity summer samples. The representative MAGs from Ia.5 and Ia.6 had slightly higher CoPTR values in the medium compared to the high salinity sample, but the CoPTR value of all others (II, IIIa.4 and V) was higher in high versus medium salinity samples. Lower growth rates did not correspond to either increases in phosphate or nitrogen transporters nor phosphate or nitrogen compound concentrations. These results suggest that there are other factors besides nitrogen and phosphorous limiting growth in these samples. Increased PHA storage transcripts in medium versus high salinities did correlate to differences in growth rates in II and IIIa.4, suggesting an imbalance of carbon to nitrogen and phosphate may be responsible for these differences in those genomospecies.

As mentioned in the main text, we observed global gene regulation patterns between samples taken at night from mid-salinity summer Delaware samples compared to all other summer Delaware samples taken from mid or high salinities, within 24-36 hours (**Fig. S5**). Several patterns were observed in COG categories D, F, J, K, L, M and O, involved in transcription, translation, replication, and cell division when comparing gene expression differences (**Fig. S10, see SI at 10.6084/m9.figshare.19701484**). Representative MAGs all had increases in gene expression from these COG categories in the mid-salinity day and high salinity samples compared to the medium night salinity samples. Expression of genes within COG categories E|G, E|T, G, I, Q and V were generally opposite, where the medium night salinity samples had increased expression compared to the others, except for *pha*C in genomospecies from II and IIIa.4. However, gene expression patterns within other COG categories (C, E, P) were mixed. Expression of genes involved in phosphate and ammonium transport, along with the energy conservation cytochrome genes *atp*ABCDG, glutamine and glutamate synthetases (*gln*A, *glt*BD) and isoleucine biosynthesis (*leu*C), among others, were increased in the high salinity and medium day salinity samples compared to the medium night salinity samples. As mentioned in the main text, the observed growth rates in representative MAGs (**Table S2**) were opposite to this and paralleled the increased concentrations of ammonium and phosphate in the mid night sample compared to the rest (**Figs. S5, S10**).

The common gene expression patterns seen in this study generally reflect differences in salinity, amino acid metabolism, and inorganic ion transport. However, like that of the comparison between the bay summer samples in the main text, expression of genes seemed to be globally regulated across all representative genomospecies, with transcripts of genes involved in replication, transcription and translation were typically increased in the Chesapeake spring high compared to medium salinities but increased in Chesapeake summer medium compared to high salinities. Genes in COG category O (Post-translational modification, protein turnover, chaperone functions) were generally increased in the opposite condition in the Chesapeake summer medium versus high salinity samples but not in the other comparisons, where the patterns were mixed. Additionally, we did not observe many common patterns in gene expression from COG category G (carbohydrate metabolism/transport), rather they were more genomospecies specific (**see SI at 10.6084/m9.figshare.19701484**). Overall, differences in observed gene expression patterns may associate with changes in either organic matter content or growth stage, as observed in other marine bacteria (43, 44). In entirety, our results suggest both a common regulation of certain pathways when the Pelagibacterales encounters a certain condition, but also a mixed regulation of others, especially those in carbohydrate transport and metabolism, that reflects individual genetic repertoires and niche space of these SAR11 MAGs representative of different genomospecies.
