## Supplementary material for "Controls of SAR11 subclade abundance, diversity, and growth in two Mid-Atlantic estuaries": Table S1

**Table S1.** Properties of SAR11 MAGs, SAGs and genomes used in this study. Subclades were defined based on identities to previously characterized MAGs, SAGs and genomes via phylogenomics and AAI/ANI analyses. Genome/MAG properties were determined with Anvi’o.

| **MAG/genome** | **NCBI biosample**  **accession#** | **Sub-**  **clade** | **CoPTR**  **rep.** | **DESeq2** | **95% rep.** | **total length (bp)** | **#contigs** | **N50** | **GC content** | **% completion** | **% redundancy** |
| --- | --- | --- | --- | --- | --- | --- | --- | --- | --- | --- | --- |
| AG_414_M23 | SAMN08886402 | Ib.3 |  |  | yes | 1104608 | 38 | 63912 | 28.82 | 94.37 | 0 |
| AG_430_F16 | SAMN08886473 | Ib.4 |  |  | yes | 1108048 | 13 | 264091 | 29.56 | 70.42 | 1.41 |
| AG_464_P07 | SAMN08886571 | Ic |  |  | yes | 1138562 | 46 | 77204 | 29.71 | 81.69 | 0 |
| ARS1026 | SAMN07619614 | Ic |  |  |  | 983187 | 35 | 45759 | 30.57 | 73.24 | 4.23 |
| BACL1_MAG_120920_64 | SAMN03742009 | Ia.1 |  |  |  | 711101 | 196 | 3754 | 30.78 | 47.89 | 0 |
| BACL5_MAG_120705_12 | SAMN03741994 | Ia.5 |  |  |  | 1056063 | 155 | 7115 | 31.90 | 77.46 | 1.41 |
| BACL5_MAG_120813_20 | SAMN03741995 | Ia.5 |  |  |  | 667156 | 170 | 4184 | 30.53 | 71.83 | 0 |
| BACL5_MAG_120820_39 | SAMN03741996 | Ia.5 |  |  |  | 1048735 | 154 | 9819 | 29.16 | 87.32 | 0 |
| BACL5_MAG_121015_10 | SAMN03741997 | Ia.5 |  |  |  | 1095111 | 189 | 6658 | 29.48 | 77.46 | 0 |
| BACL5_MAG_121128_54 | SAMN03741998 | Ia.5 |  |  |  | 922610 | 175 | 6291 | 29.56 | 88.73 | 1.41 |
| CP_Spr15G08_27 | SAMN17495979 | IIIa.4 |  |  |  | 1052580 | 67 | 28096 | 30.52 | 94.36 | 0 |
| CP_Spr15G08_4 | SAMN12628525 | Ia.1 |  |  |  | 1073401 | 69 | 23672 | 29.97 | 97.18 | 0 |
| CP_Spr15L08_1 | SAMN12618548 | IIIa.2 | IIIa.2 |  | yes | 1119417 | 24 | 73217 | 31.96 | 97.18 | 1.41 |
| CP_Spr15L08_31 | SAMN12618547 | IIIa.5 |  |  | yes | 1023108 | 45 | 42723 | 31.16 | 95.77 | 1.41 |
| CP_Spr15L08_49 | SAMN17495998 | II |  |  |  | 888897 | 74 | 17918 | 30.19 | 88.73 | 0 |
| CP_Spr15L08_56 | SAMN17914822 | IIIa.5 |  |  |  | 1020527 | 51 | 42584 | 31.69 | 97.18 | 1.41 |
| CP_Spr30L08_35 | SAMN17496045 | IIIa.2 |  |  |  | 971596 | 84 | 18067 | 32.08 | 94.36 | 2.81 |
| CP_Spr30L08_58 | SAMN17496050 | IIIa.4 |  |  |  | 1077603 | 74 | 25182 | 30.74 | 97.18 | 1.40 |
| CP_Sum15G08_27 | SAMN12618233 | II |  |  |  | 998856 | 82 | 17711 | 30.11 | 91.55 | 0 |
| CP_Sum15G08_40 | SAMN12618234 | Ia.6 |  |  |  | 918225 | 82 | 17815 | 29.48 | 87.32 | 0 |
| CP_Sum15G08_43 | SAMN17496081 | IIIa.4 |  |  |  | 1021307 | 53 | 34804 | 31.41 | 95.77 | 0 |
| CP_Sum15G08_5 | SAMN12618232 | V | V | V | yes | 1234213 | 72 | 25397 | 32.90 | 87.32 | 0 |
| CP_Sum15L08_53 | SAMN17496091 | IIIa.4 |  |  |  | 991409 | 65 | 28185 | 31.31 | 90.14 | 0 |
| CP_Sum15L08_55 | SAMN12628524 | IIIa.5 |  |  | yes | 786631 | 122 | 7444 | 30.33 | 84.51 | 0 |
| CP_Sum15L08_77 | SAMN17496098 | Ia.6 | Ia.6 | Ia.6 | yes | 895354 | 77 | 19475 | 29.37 | 85.92 | 0 |
| CP_Sum20G08_2 | SAMN12628523 | IIIa.1 |  |  | yes | 964383 | 99 | 13136 | 30.64 | 81.69 | 1.41 |
| CP_Sum20G08_35 | SAMN17496103 | IIIa.4 |  |  |  | 1116755 | 60 | 32416 | 30.98 | 100.00 | 0 |
| CP_Sum20G08_55 | SAMN17496104 | II |  |  |  | 941438 | 69 | 19940 | 30.06 | 97.18 | 0 |
| CP_Sum27G08_38 | SAMN17496111 | II |  |  |  | 947902 | 85 | 16846 | 30.21 | 83.09 | 0 |
| CP_Sum27L08_15 | SAMN12628522 | IIIa.4 | IIIa.4 | IIIa | yes | 1094164 | 67 | 35486 | 31.19 | 91.55 | 0 |
| DE_Fall30L08_8 | SAMN17914823 | Ia.1 |  |  |  | 937253 | 182 | 5974 | 29.96 | 83.10 | 0 |
| DE_Spr20L08_1 | SAMN17914824 | IIIa.2 |  |  |  | 963531 | 130 | 10060 | 29.86 | 87.32 | 0 |
| DE_Spr20L08_1_2 | SAMN17914824 | Ia.1 |  |  |  | 1079511 | 33 | 51752 | 30.05 | 95.77 | 0 |
| DE_Spr20L08_17 | SAMN12628526 | IIIa.2 |  |  |  | 1062996 | 84 | 19113 | 31.52 | 90.14 | 0 |
| DE_Spr30G08_37 | SAMN17914825 | Ia.1 |  |  |  | 836479 | 125 | 8781 | 30.09 | 80.28 | 0 |
| DE_Spr30L08_1 | SAMN12628528 | Ia.1 | Ia.1 | Ia.1 | yes | 1066630 | 89 | 22675 | 29.95 | 95.77 | 0 |
| DE_Sum22DL08_11 | SAMN17914826 | IIIa.4 |  |  |  | 928580 | 170 | 6912 | 31.13 | 80.28 | 0 |
| DE_Sum22DL08_27 | SAMN17496223 | IIIa.4 |  |  |  | 1007645 | 32 | 58945 | 31.00 | 94.36 | 0 |
| DE_Sum22DL08_33 | SAMN12618555 | IIIa.4 |  |  |  | 935134 | 30 | 55832 | 31.27 | 88.73 | 0 |
| DE_Sum22DL08_48 | SAMN17914827 | Ia.1 |  |  |  | 1032240 | 82 | 23541 | 30.05 | 85.92 | 0 |
| DE_Sum22DL08_52 | SAMN17914828 | Ia.5 | Ia.5 | Ia.5 | yes | 983531 | 54 | 38048 | 30.01 | 84.51 | 0 |
| DE_Sum22DL08_73 | SAMN12618556 | V |  |  |  | 1087288 | 26 | 84544 | 32.32 | 78.87 | 0 |
| DE_Sum22DL08_in_19 | SAMN25815057 | II |  |  |  | 1020477 | 78 | 20813 | 30.27 | 95.77 | 0 |
| DE_Sum29DG08_42 | SAMN12628527 | Ia.6 |  |  |  | 876443 | 69 | 23829 | 29.34 | 76.06 | 1.41 |
| DE_Sum29DG08_44 | SAMN17914829 | II |  |  |  | 889835 | 91 | 13439 | 30.36 | 88.73 | 0 |
| DE_Sum29DL08_25 | SAMN17914830 | Ia.3.VIII.other | Ia.3 | Ia.3 | yes | 1156098 | 173 | 8650 | 29.78 | 84.51 | 2.82 |
| DE_Sum29DL08_3 | SAMN17914831 | Ia.6 |  |  |  | 1048190 | 159 | 9485 | 30.19 | 78.87 | 0 |
| DE_Sum29DL08_37 | SAMN17914832 | Ia.1 |  |  |  | 818446 | 72 | 14349 | 29.57 | 85.92 | 0 |
| DE_Sum29DL08_4 | SAMN17914833 | Ia.3.VIII.other |  |  |  | 1018566 | 158 | 8055 | 29.61 | 83.10 | 2.82 |
| DE_Sum29DL08_52 | SAMN17914834 | II | II | II | yes | 1003485 | 80 | 23353 | 30.41 | 94.37 | 2.82 |
| DE_Sum29NG08_1 | SAMN17914835 | II |  |  |  | 765322 | 81 | 13929 | 29.40 | 71.83 | 0 |
| DE_Sum29NG08_65 | SAMN17914836 | Ia.6 |  | Ia.6 |  | 935939 | 91 | 16383 | 29.47 | 85.92 | 0 |
| DE_Sum29NL08_145 | SAMN17914837 | Ia.1 |  |  |  | 949513 | 79 | 17130 | 30.06 | 84.51 | 0 |
| DE_Sum29NL08_47 | SAMN12628529 | II |  |  |  | 902525 | 65 | 29721 | 30.18 | 85.92 | 0 |
| FZCC0015 | SAMN09644087 | Ia.3.V_VI_VII |  |  | yes | 1364101 | 1 | 1364101 | 29.20 | 98.59 | 0 |
| GOM_A1 | SAMN03944292 | Ib.1 |  |  | yes | 1285855 | 37 | 84415 | 29.38 | 92.96 | 0 |
| HIMB058 | SAMN02440920 | IIa.B |  |  | yes | 1100317 | 48 | 38257 | 29.56 | 95.77 | 0 |
| HIMB083 | SAMN02597166 | Ia.3.V_VI_VII |  |  | yes | 1395997 | 1 | 1395997 | 29.16 | 98.59 | 0 |
| HIMB122 | SAMN02744629 | Ia.3.V_VI_VII |  |  | yes | 1453515 | 1 | 1453515 | 29.29 | 97.18 | 0 |
| HIMB1321 | SAMN02744631 | Ia.6 |  |  | yes | 1320749 | 1 | 1320749 | 29.03 | 98.59 | 0 |
| HIMB140 | SAMN03402370 | Ia.3.V_VI_VII |  |  | yes | 1437930 | 1 | 1437930 | 29.37 | 98.59 | 0 |
| HIMB4 | IMG# 2503754001 | Ia.6 |  |  |  | 1382291 | 1 | 1382291 | 28.98 | 98.59 | 0 |
| HIMB5 | SAMN00016662 | Ia.6 |  |  | yes | 1343202 | 1 | 1343202 | 28.65 | 98.59 | 0 |
| HIMB59 | SAMN00010387 | V |  |  | yes | 1410127 | 1 | 1410127 | 32.24 | 97.18 | 0 |
| HTCC1002 | SAMN02436088 | Ia.1 |  |  |  | 1326048 | 2 | 1324008 | 29.96 | 98.59 | 0 |
| HTCC1013 | SAMN02441456 | Ia.1 |  |  |  | 1299699 | 3 | 741014 | 29.78 | 98.59 | 0 |
| HTCC1016 | SAMN02256429 | Ia.1 |  |  |  | 1297098 | 4 | 794419 | 29.73 | 97.18 | 0 |
| HTCC1040 | SAMN02256395 | Ia.1 |  |  |  | 1274624 | 1 | 1274624 | 29.58 | 98.59 | 0 |
| HTCC1062 | SAMN02603690 | Ia.1 |  |  |  | 1308759 | 1 | 1308759 | 29.61 | 98.59 | 0 |
| HTCC7211 | SAMN02436224 | Ia.3.I.IV |  |  | yes | 1456888 | 1 | 1456888 | 29.01 | 98.59 | 0 |
| HTCC7214 | SAMN02841172 | Ia.3.I.IV |  |  | yes | 1375060 | 1 | 1375060 | 29.24 | 97.18 | 0 |
| HTCC7217 | SAMN02841150 | Ia.3.I.IV |  |  | yes | 1433611 | 1 | 1433611 | 28.81 | 98.59 | 0 |
| HTCC9022 | SAMN02440781 | Ia.3.VIII.other |  |  |  | 1355934 | 3 | 805404 | 29.59 | 98.59 | 0 |
| HTCC9565 | SAMN03402630 | Ia.1 |  |  | yes | 1279674 | 3 | 763172 | 28.95 | 97.18 | 0 |
| IMCC9063 | SAMN02603337 | IIIa.2 |  |  | yes | 1284727 | 1 | 1284727 | 31.64 | 98.59 | 1.41 |
| LSUCC0530 | SAMN07786750 | IIIb |  |  | yes | 1160202 | 1 | 1160202 | 29.01 | 97.18 | 0 |
| MED_G40 | SAMN06890651 | IIa.B |  |  | yes | 911758 | 8 | 205872 | 29.66 | 83.10 | 0 |
| MED_G43 | SAMN06890654 | IIIa.1 |  |  |  | 421449 | 22 | 18425 | 29.31 | 63.38 | 0 |
| MED104 | SAMN07619616 | IIa |  |  |  | 868752 | 73 | 12239 | 29.61 | 77.46 | 15.49 |
| MED727 | SAMN07619622 | IIa |  |  |  | 503620 | 23 | 32957 | 29.73 | 74.65 | 0 |
| MED769 | SAMN07619623 | Ib.5 |  |  | yes | 1263754 | 48 | 30066 | 28.74 | 83.10 | 8.45 |
| MED817 | SAMN07619625 | IIa |  |  |  | 775528 | 61 | 13397 | 28.56 | 70.42 | 0 |
| MED827 | SAMN07619645 | IIa |  |  |  | 857302 | 26 | 50206 | 28.69 | 54.93 | 2.82 |
| MED832 | SAMN07619646 | IIIa.3 |  |  |  | 958664 | 14 | 135228 | 29.59 | 50.70 | 5.63 |
| NORP132 | SAMN07568946 | Ib.6 |  |  |  | 978135 | 107 | 11970 | 30.96 | 73.24 | 2.82 |
| NORP187 | SAMN07569001 | Ic |  |  |  | 848811 | 70 | 18840 | 31.64 | 39.44 | 2.82 |
| NP1 | SAMN11347529 | Ia.1 |  | NP1 | yes | 1365705 | 1 | 1365705 | 29.58 | 98.59 | 1.41 |
| QL1 | SAMN02798134 | IIIa.5 |  |  |  | 934186 | 87 | 19764 | 31.86 | 77.46 | 14.08 |
| RS39 | SAMN06562974 | Ia.4 |  |  | yes | 1200090 | 1 | 1200090 | 29.24 | 97.18 | 0 |
| RS40 | SAMN06562975 | Ib.1 |  |  | yes | 1378905 | 1 | 1378905 | 29.41 | 97.18 | 0 |
| SAG_MED01 | SAMN15685163 | Ia.4 |  |  | yes | 1167589 | 19 | 154766 | 28.22 | 95.77 | 0 |
| SAG_MED05 | SAMN15685167 | IIa |  |  | yes | 955906 | 25 | 101374 | 29.45 | 85.92 | 0 |
| SAG_MED06 | SAMN15685168 | Ia.3.VIII.other |  |  |  | 1203374 | 29 | 85407 | 29.17 | 88.73 | 0 |
| SAG_MED08 | SAMN15685170 | Ia.3.V_VI_VII |  |  | yes | 1092371 | 31 | 95578 | 29.68 | 88.73 | 1.41 |
| SAG_MED11 | SAMN15685173 | Ia.4 |  |  | yes | 1090219 | 16 | 240570 | 28.82 | 98.59 | 0 |
| SAG_MED12 | SAMN15685174 | Ia.4 |  |  |  | 919059 | 28 | 66544 | 28.84 | 84.51 | 0 |
| SAG_MED13 | SAMN15685175 | Ia.4 |  |  | yes | 991550 | 32 | 59163 | 28.59 | 87.32 | 0 |
| SAG_MED15 | SAMN15685177 | Ib.5 |  |  | yes | 1189410 | 17 | 199546 | 28.91 | 92.96 | 0 |
| SAG_MED16 | SAMN15685178 | Ib.2 |  |  |  | 869866 | 44 | 34855 | 29.71 | 74.65 | 1.41 |
| SAG_MED19 | SAMN15685181 | Ia.4 |  |  | yes | 990483 | 34 | 69724 | 28.33 | 90.14 | 0 |
| SAG_MED22 | SAMN15685184 | Ia.3.V_VI_VII |  |  | yes | 1175045 | 14 | 175820 | 29.46 | 94.37 | 0 |
| SAG_MED28 | SAMN15685189 | IIa |  |  | yes | 967204 | 23 | 87518 | 29.56 | 91.55 | 1.41 |
| SAG_MED29 | SAMN15685190 | IIa |  |  | yes | 925266 | 28 | 95465 | 28.90 | 78.87 | 0 |
| SAG_MED33 | SAMN15685194 | Ia.3.V_VI_VII |  |  |  | 1065163 | 19 | 98442 | 29.78 | 92.96 | 0 |
| SAG_MED37 | SAMN15685197 | Ia.4 |  |  | yes | 949129 | 35 | 102181 | 28.99 | 78.87 | 1.41 |
| SAG_MED47 | SAMN15685205 | Ia.4 |  |  | yes | 1113380 | 29 | 104783 | 28.56 | 91.55 | 1.41 |
| SAG_MED48 | SAMN15685206 | Ia.3.I.IV |  |  |  | 1037700 | 40 | 59913 | 29.34 | 57.75 | 0 |
| SAG_MED49 | SAMN15685207 | Ia.3.I.IV |  |  |  | 1157512 | 30 | 105173 | 29.03 | 88.73 | 0 |
| SAG_MED50 | SAMN15685208 | Ia.3.VIII.other |  |  | yes | 1231880 | 35 | 67996 | 29.31 | 85.92 | 0 |
| SCGC_AAA795_A20 | SAMN02256429 | Ia.3.V_VI_VII |  |  |  | 1114625 | 31 | 106301 | 29.17 | 87.32 | 0 |
| SCGC_AAA795_B16 | SAMN02256395 | Ib.1 |  |  |  | 497805 | 40 | 18649 | 29.27 | 60.56 | 0 |
| SCGC_AAA795_D22 | SAMN02603690 | Ia.4 |  |  | yes | 978204 | 33 | 79594 | 29.16 | 76.06 | 0 |
| SCGC_AAA795_E07 | SAMN02436224 | IIa.B |  |  |  | 613294 | 27 | 50970 | 29.72 | 71.83 | 0 |
| SCGC_AAA795_E22 | SAMN02841172 | Ib.5 |  |  |  | 775868 | 34 | 56618 | 29.29 | 74.65 | 4.23 |
| SCGC_AAA795_F16 | SAMN02841150 | Ib.4 |  |  | yes | 895136 | 22 | 101331 | 28.86 | 84.51 | 0 |
| SCGC_AAA795_J21 | SAMN02440710 | Ia.3.V_VI_VII |  |  |  | 825765 | 23 | 120490 | 29.00 | 67.61 | 0 |
| SCGC_AAA795_M18 | SAMN02440781 | Ib.1 |  |  |  | 1029034 | 37 | 66910 | 29.10 | 71.83 | 0 |
| SCGC_AAA795_M22 | SAMN03402630 | Ib.4 |  |  | yes | 811889 | 27 | 66137 | 29.38 | 85.92 | 0 |
| SCGC_AAA795_O19 | SAMN02603337 | Ia.3.V_VI_VII |  |  |  | 776039 | 27 | 70048 | 29.06 | 74.65 | 0 |
| SCGC_AAA795_O20 | SAMN07786750 | Ia.4 |  |  | yes | 985052 | 36 | 81405 | 28.89 | 78.87 | 0 |
| SCGC_AAA795_P11 | SAMN07619616 | Ia.3.V_VI_VII |  |  |  | 914706 | 52 | 39306 | 29.36 | 78.87 | 0 |
| SCGC_AAA797_I19 | SAMN07619622 | Ia.3.V_VI_VII |  |  |  | 972854 | 23 | 157078 | 28.88 | 77.46 | 0 |
| UBA2154 | SAMN07619623 | Ic |  |  | yes | 1076758 | 126 | 11153 | 31.10 | 74.65 | 1.41 |
| WB5_1B_032_PEL3 | SAMN10222973 | IIIb |  |  |  | 395961 | 97 | 4140 | 29.60 | 59.15 | 0 |
| WB5_1B_059_PEL4 | SAMN10223001 | IIIb |  |  |  | 598122 | 118 | 6089 | 29.26 | 70.42 | 0 |
| WB7_2A_005_PEL5 | SAMN10223253 | IIIb |  |  |  | 794547 | 178 | 5017 | 29.20 | 64.79 | 0 |
| WB7_2xF_001_PEL2 | SAMN10223262 | IIIb |  |  |  | 596320 | 104 | 7019 | 29.98 | 74.65 | 0 |
| WB7_3xA_007_PEL7 | SAMN10223280 | IIIa.4 |  |  |  | 635268 | 139 | 4930 | 32.06 | 56.34 | 0 |
| WB7_3xF_003_PEL6 | SAMN10223271 | Ia.5 |  |  |  | 686055 | 143 | 5520 | 30.47 | 52.11 | 0 |
| WB8_4xD_006_PEL8 | SAMN10223526 | Ia.6 |  |  |  | 612627 | 128 | 5485 | 29.89 | 60.56 | 0 |
| WB8_6_001_PEL1 | SAMN10223538 | IIIb |  |  | yes | 978845 | 153 | 8071 | 35.99 | 83.10 | 0 |
