## Supplementary material for "Controls of SAR11 subclade abundance, diversity, and growth in two Mid-Atlantic estuaries": Table S2

| **Metagenome** | **Ia.1** | **Ia.3** | **Ia.5** | **Ia.6** | **II** | **IIIa.2** | **IIIa.4** | **V** | **Average** |
| --- | --- | --- | --- | --- | --- | --- | --- | --- | --- |
| CP_Spr15 | 0.53 |  | 0.76 | 0.37 | 0.63 | 0.55 | 0.61 |  | 0.58 |
| CP_Spr31 | 0.60 | 0.67 | 0.86 | 0.39 | 0.77 | 0.40 | 0.71 |  | 0.63 |
| CP_Sum15 |  |  | 1.23 | 0.37 | 0.54 |  | 0.51 | 0.70 | 0.67 |
| CP_Sum27 | 0.44 | 0.37 | 1.10 | 0.34 | 0.55 |  | 0.56 | 0.78 | 0.59 |
| DE_Fall15 | 0.37 | 1.13 | 0.94 | 1.23 | 1.24 |  | 0.62 | 1.59 | 1.02 |
| DE_Fall30 | 0.55 | 0.51 | 0.68 | 0.45 | 0.67 |  | 0.54 | 0.82 | 0.60 |
| DE_Spr20 | 0.57 |  |  |  | 0.56 | 0.47 |  |  | 0.53 |
| DE_Spr30 | 0.66 | 1.05 |  |  |  | 0.39 |  |  | 0.70 |
| DE_Sum22N | 0.40 | 1.00 | 0.76 | 1.11 | 1.30 |  | 0.46 | 1.41 | 0.92 |
| DE_Sum22D | 0.43 | 1.01 | 0.72 | 1.26 | 1.08 |  | 0.54 | 1.11 | 0.88 |
| DE_Sum29N | 0.52 | 0.92 | 0.74 | 1.14 | 1.11 | 0.36 | 0.55 | 1.06 | 0.80 |
| DE_Sum29D | 0.46 | 0.77 | 0.72 | 0.87 | 0.95 |  |  | 1.11 | 0.81 |
| **Average** | 0.50 | 0.83 | 0.85 | 0.75 | 0.86 | 0.43 | 0.57 | 1.07 |  |

**Table S2.** CoPTR values from a representative MAG in the indicated subclade from Delaware (DE) and Chesapeake (CP) Bay metagenomes from the less than 0.8 µm size fraction. The number after Spr (Spring), Sum (Summer or Fall is the salinity in PSU. N = night, D = day. Representative MAGs are listed in **Table S1**.
