## Supplementary figures and images for "Controls of SAR11 subclade abundance, diversity, and growth in two Mid-Atlantic estuaries"

### Fig S3

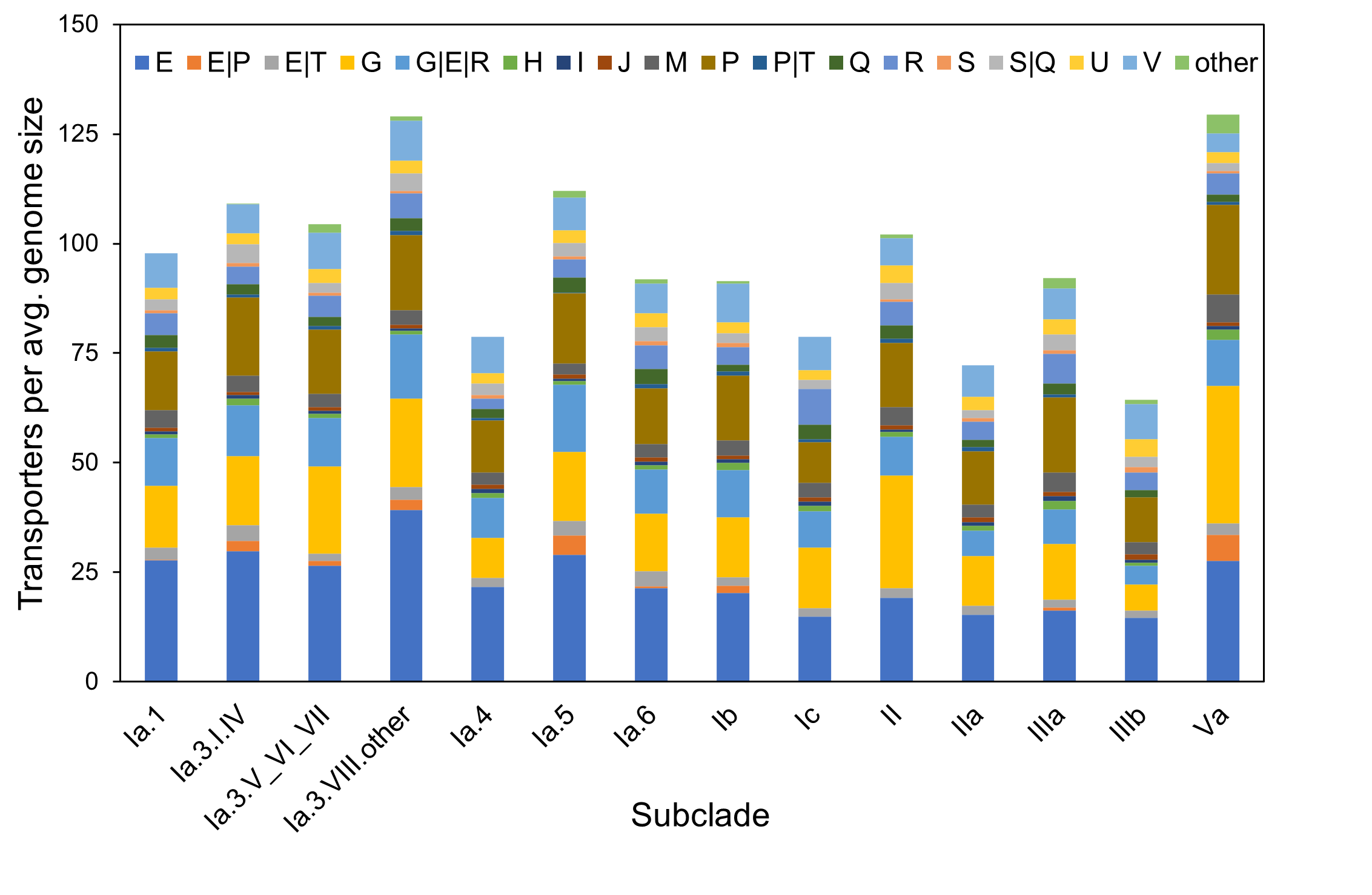

### Fig S4

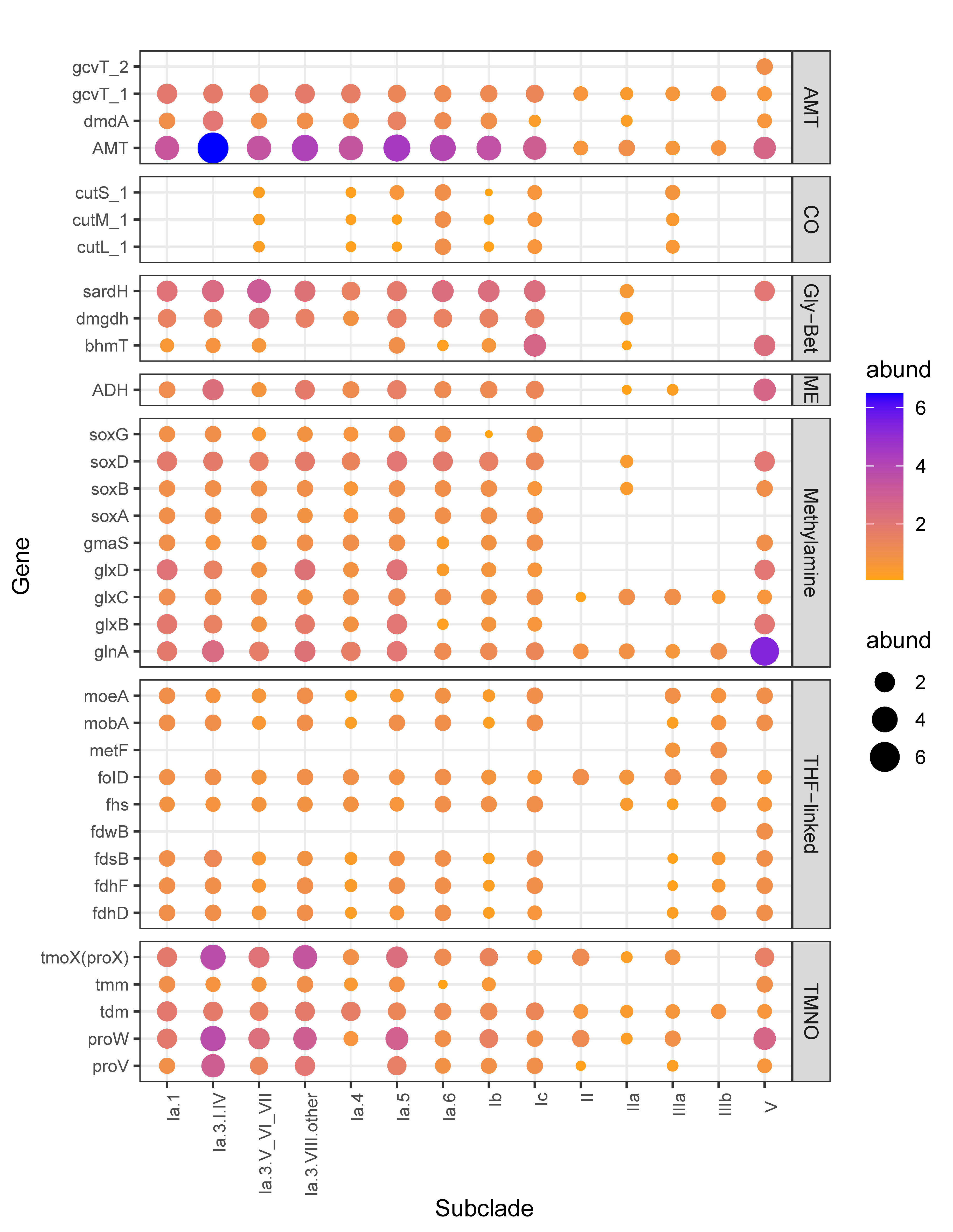

### Fig S5

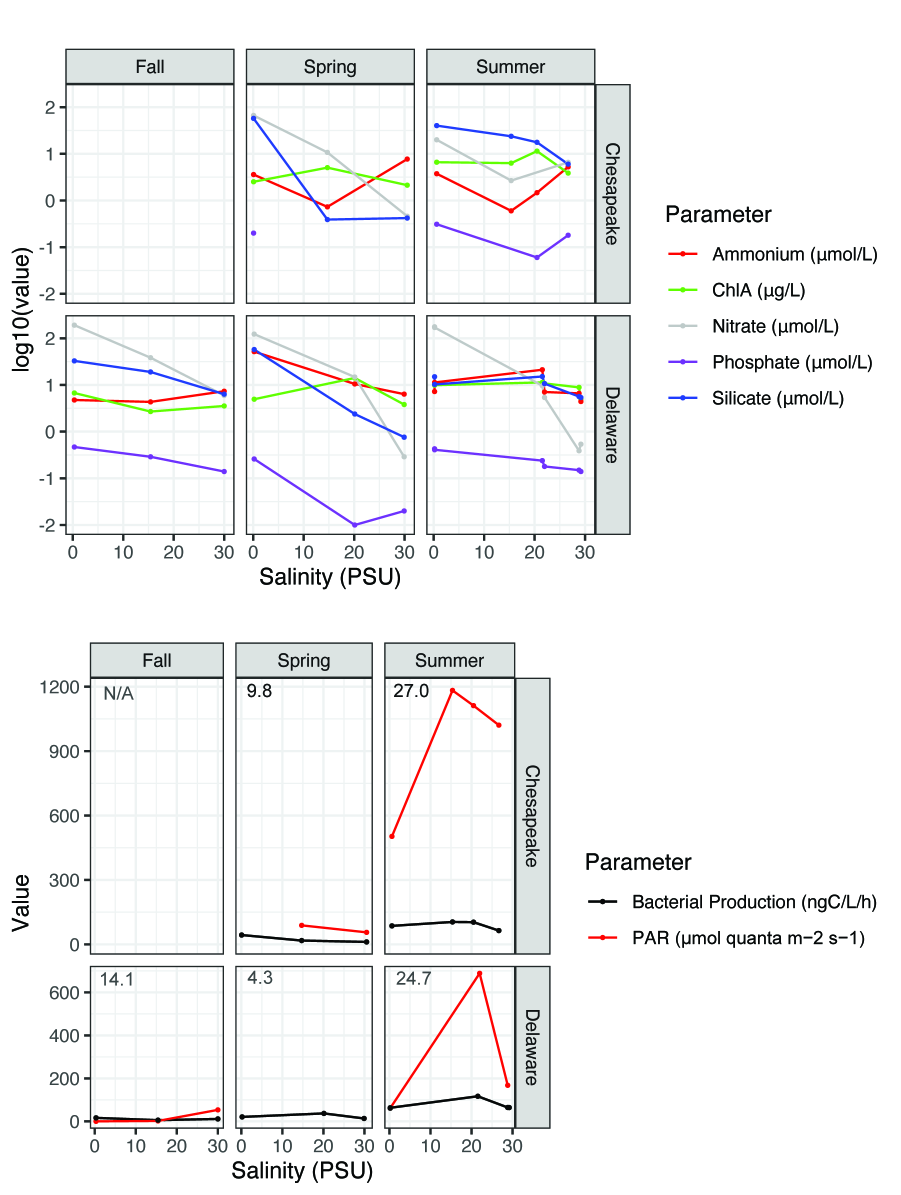

### Fig S6

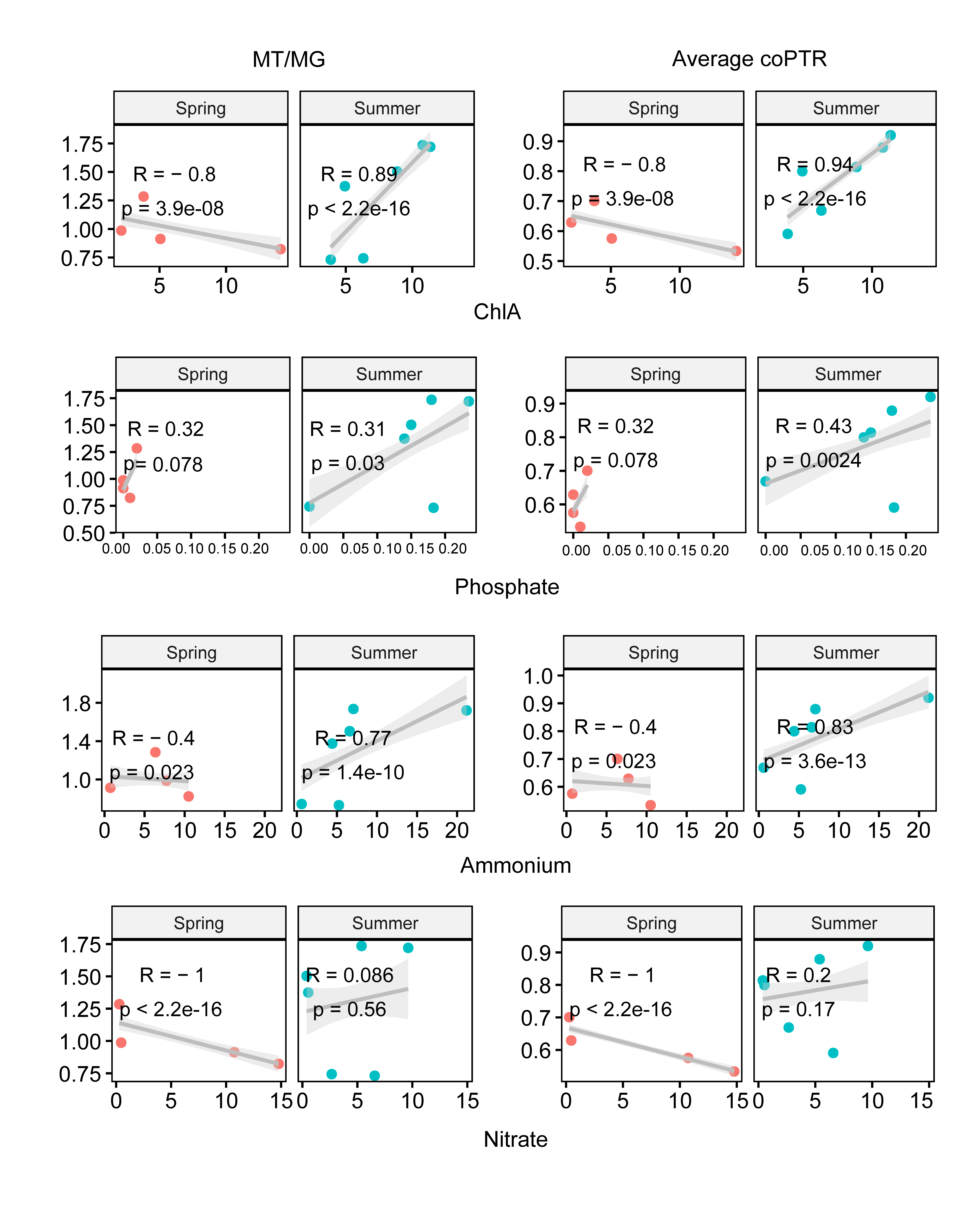

### Fig S10

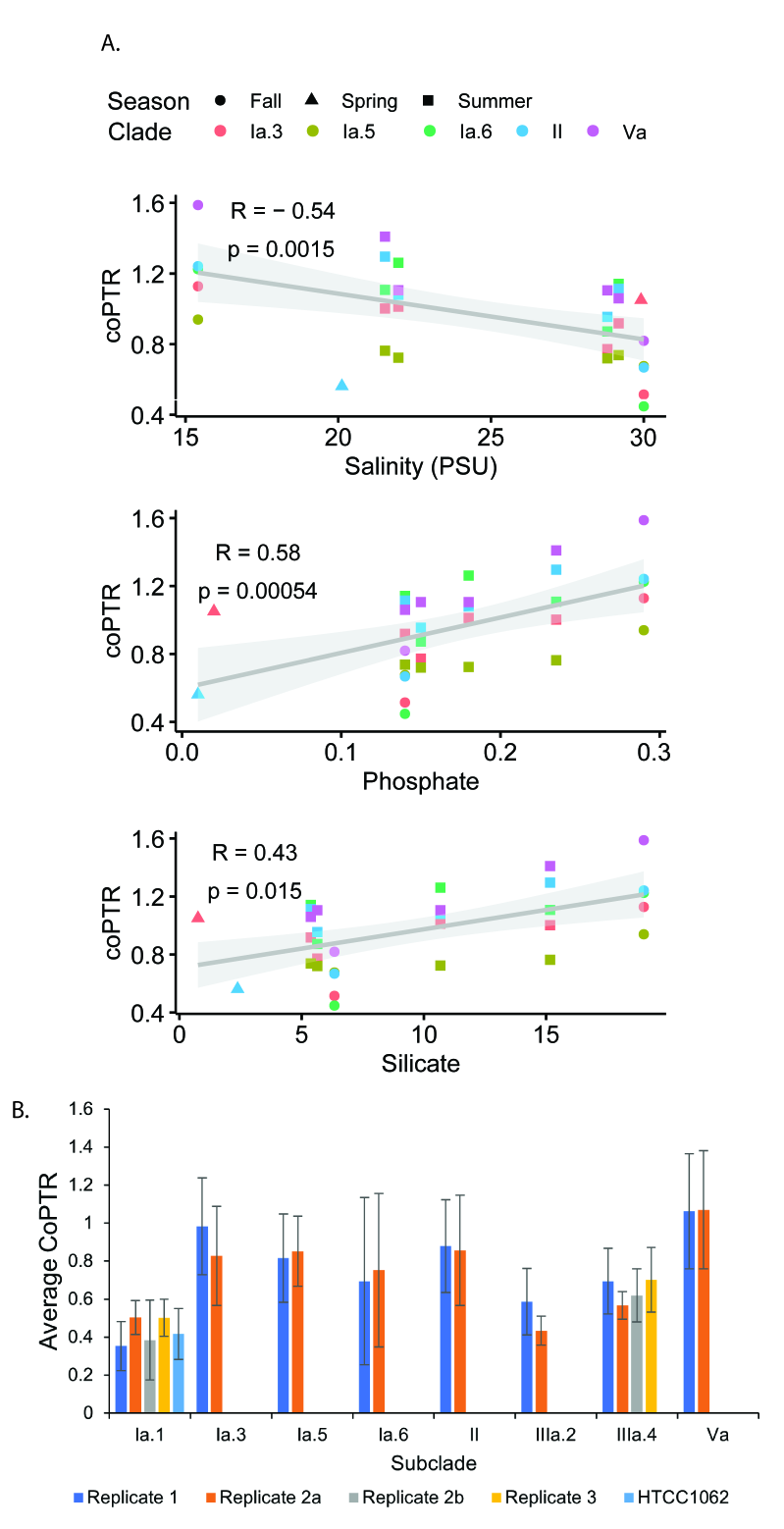

### Fig S10

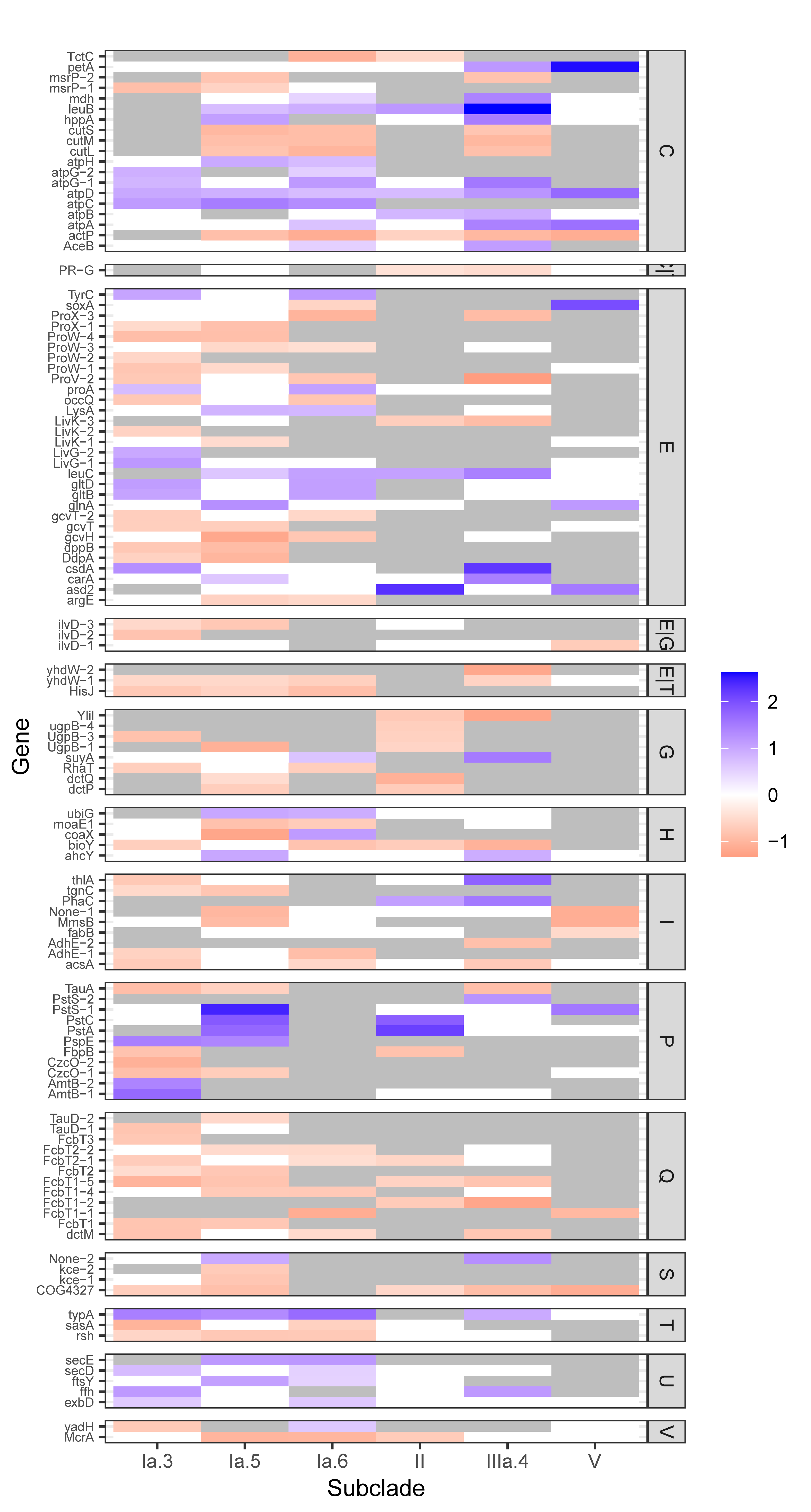
